# Microbial colonization of tannin-rich tropical plants: interplay between degradability, methane production and tannin disappearance in the rumen

**DOI:** 10.1101/2021.08.12.456105

**Authors:** Moufida Rira, Diego P. Morgavi, Milka Popova, Gaёlle Maxin, Michel Doreau

**Affiliations:** INRAE, VetAgro Sup, UMR1213 Herbivores, F-63122 — Saint-Genès-Champanelle, France; Ecole Nationale Supérieure de Biotechnologie, Ali Mendjli, BP E66, 25100 — Constantine, Algeria

**Keywords:** tropical plants, hydrolysable tannins, condensed tannins, methane, in situ degradability, rumen microbes

## Abstract

Condensed tannins in plants are found free and attached to protein and fibre but it is not known whether these fractions influence rumen degradation and microbial colonization. This study explored the rumen degradation of tropical tannins-rich plants and the relationship between condensed tannins fractions’ disappearance and microbial communities colonising plant particles. Leaves from *Calliandra calothyrsus, Gliricidia sepium*, and *Leucaena leucocephala*, pods from *Acacia nilotica* and the leaves of two agricultural by-products: Manihot esculenta and *Musa* spp. were incubated in situ in the rumen of dairy cows. An in vitro approach was also used to assess the effects of these plants on rumen fermentation parameters. All plants contained more than 100 g/kg of condensed tannins with a large proportion (32 to 61%) bound to proteins. *Calliandra calothyrsus* had the highest concentration of condensed tannins at 361 g/kg, whereas *Acacia nilotica* was particularly rich in hydrolysable tannins (350 g/kg). Hydrolysable and free condensed tannins from all plants completely disappeared after 24 h incubation in the rumen. Disappearance of protein-bound condensed tannins was variable with values ranging from 93% for *Gliricidia sepium* to 21% for *Acacia nilotica*. In contrast, fibre-bound condensed tannins disappearance averaged ~82% and did not vary between plants. Disappearance of bound fractions of condensed tannins was not associated with degradability of plant fractions. The presence of tannins interfered with the microbial colonisation of plants. Each plant had distinct bacterial and archaeal communities after 3 and 12 h of incubation in the rumen and distinct protozoal communities at 3 h. Adherent communities in tannin-rich plants had a lower relative abundance of fibrolytic microbes, notably *Fibrobacter* spp. Whereas, archaea diversity was reduced in high tannin-containing *Calliandra calothyrsus* and *Acacia nilotica* at 12 h of incubation. Concurrently, in vitro methane production was lower for *Calliandra calothyrsus, Acacia nilotica* and *Leucaena leucocephala* although for the latter total volatile fatty acids production was not affected and was similar to control. Here we show that the total amount of hydrolysable and condensed tannins contained in a plant govern the interaction with rumen microbes affecting degradability and fermentation. The effect of protein- and fibre-bound condensed tannins on degradability is less important.

## Introduction

Feed availability is a major limitation in many tropical ruminant production systems. One way for farmers to increase forage availability in these grazing systems is to use leguminous trees for supplementing diets. Leaves from leguminous trees are a high-protein feed resource that balances the nutrition of ruminants grazing tropical grasses poor in protein (Roothaert and Paterson, 1997). However, they remain underutilised because of the presence of tannins that are often found at high concentrations. High concentrations of tannins can reduce voluntary feed intake and nutrient digestibility (Frutos et al., 2004). The decrease in nutritive value is associated with tannins’ property to bind to proteins, both from the diet and digestive enzymes. Tannins also bind to structural carbohydrates present in plant cell walls (Mueller-Harvey et al., 2019). Despite these adverse effects, a low concentration of tannins in the diet can improve nitrogen (N) utilisation efficiency and has a positive enteric methane-reduction effect (Goel and Makkar, 2012). This latter effect is because tannins may reduce organic matter digestibility in the rumen even when total-tract digestibility is unchanged, or because they inhibit microbial populations, or both (Frutos et al., 2004).

Tannins are conventionally classified into two major groups: hydrolysable (HT) and condensed tannins (CT). Hydrolysable tannins consist of polyphenols (gallic acid and/or hexahydroxydiphenic acid) ester-linked to a hexose moiety. They are categorised according to their structural characteristics into two subgroups: gallotannins and ellagitannins. In contrast, CT are polymers of varying molecular weight composed of flavan-3-ol (e.g., catechin) or flavan-3,4-diol (proanthocyanidins) linked by C–C or C–O–C bonds. Condensed tannins are found in different fractions in plants: free, protein-bound, and fibre-bound (Mueller-Harvey and McAllan, 1992, Terrill et al., 1992, Schofield et al., 2001). A better understanding of the effects of HT-rich and CT-rich forages on nutrient digestibility and methane mitigation properties would improve the management of such resources. To this end, the relationship between feed degradation and the disappearance of CT fractions must be established for developing feeding strategies overcoming undesirable effects when using tannin-rich forages. We hypothesised that rumen microbial degradation of tannin-rich forages is influenced not only by the abundance of tannins but also by their chemical form and binding to plants’ structural components. This knowledge would be of considerable importance for the efficient utilisation of these forages in the tropics.

Our objective was to study the degradation of tannin-rich forages in the rumen and establish the relationship with the amount and disappearance of their different tannin fractions. We connected these effects on forage degradation and tannin disappearance with the microbial communities colonising feed particles in the rumen and with fermentation parameters. We used six tropical forages from leguminous shrubs and agricultural by-products with differing amounts and nature of tannins. We carried out 1) an in situ experiment, in order to determine the ruminal degradation of forage components, including different tannin fractions, and the colonisation of feed particles by microbes, and 2) an in vitro experiment, in order to measure methane production and feed fermentation.

## Methods

The use of experimental animals followed the guidelines for animal research of the French Ministry of Agriculture and other applicable guidelines and regulations for animal experimentation in the European Union. Animals were housed at the INRAE UE1414 Herbipôle Unit (Saint Genès Champanelle, France; https://doi.org/10.15454/1.5572318050509348E12). Procedures were approved by French Ministry of Education and Research (APAFIS #8218-20161151782412).

### Plant material

Four browse species and two crops by-products rich in tannins and available in the tropics were selected. The browse species were leguminous shrubs: *Acacia nilotica* pods, *Calliandra calothyrsus, Gliricidia sepium* and *Leucaena leucocephala* leaves. The by-products were cassava (*Manihot esculenta*) leaves and banana (*Musa* spp.) leaves. These species are consumed by ruminants and have a large range of CT and HT content.

*Acacia nilotica* samples were collected from the Ferlo region of Senegal (15°N, 15°W) having a mean annual rainfall of 200-400 mm and a mean temperature of 28-30°C. *Gliricidia sepium, Leucaena leucocephala, Manihot esculenta* and *Musa* spp. were collected in Guadeloupe, Basse-Terre Island, France (16°N, 61°W), having a mean annual rainfall of 1500-2000 mm and a mean temperature of 24-28°C. *Calliandra calothyrsus* was collected from native shrubs in the south of the Réunion Island, France (21°S, 55°E) having a mean annual rainfall of 1000-1500 mm and a mean temperature of 20-24°C; leaves were harvested at a late vegetative stage.

Following collection, fresh material was dried at 40 °C to avoid degradation or modification of tannins, ground to pass through a 1-mm grid and placed in air-tight plastic bags. Upon reception at the laboratory in Metropolitan France, samples were stored at room temperature until use for *in situ* and *in vitro* measurements. Forages poor in tannins were used as a control when necessary for a better understanding of processes. For studying microbial colonisation of feed particles, the control forage was the natural grassland hay used for feeding animals. For studying changes in methane production and feed fermentation, the control forage was hay from a 75-d regrowth of natural grassland based on *Dichanthium* spp. harvested in Guadeloupe, Grande-Terre Island, France.

### Experiment 1: In situ rumen degradation

Three rumen-cannulated Holstein non-lactating cows were used for the study with an average body weight of 737 ± 40 kg. Cows were housed in individual pens and fed natural grassland hay, first cycle harvested in Auvergne, France. A fixed amount of 7.6 kg DM of hay was offered to each cow twice a day: 2/3 at 0900 h, 1/3 at 1600 h. Water was on free access. In situ ruminal incubations started after 15 days of adaptation to the diet.

Feeds were incubated in the rumen for 3, 6, 12, 24, 48 and 96 h. Three grams of ground samples were put into 5.5 × 12 cm polyester bags (pore size ca. 50 μm, model R1020, Ankom, Fairport, NY). Bags were hooked to a stainless-steel weight and inserted in the ventral sac of the rumen at 0800 h. Two successive series of incubations were performed for each cow. Each series had one bag per feed and incubation time for measurements of DM, N and NDF degradation, two bags per feed for measurements of CT disappearance at 24 h of incubation to get enough residues for further tannin analyses and two additional bags per feed for measurements of microbial colonisation at 3 h and at 12 h. At each designated incubation time, bags for measurements of DM, N and NDF degradation and tannin disappearance were removed from the rumen, immersed in cold water and then washed under running tap water until the water became clear. Zero-time disappearance was obtained by washing two non-incubated bags per feed as described above. Bags were kept at 4 °C for 48 h then washed in a washing machine without detergent until clean water was obtained (4 cycles of 10 min), dried at 40 °C for 96 h and weighed to determine rumen residual DM content. Bags for microbial analysis, taken out after 3 and 12 h of incubation, were gently squeezed, put on ice and immediately transferred to the lab where they were washed three times in phosphate-buffered saline at 4°C for 5 min on a rocking shaker at ~40 cycles per min. Bags were then snap-frozen in liquid N and stored at −80°C until gDNA extraction.

### Experiment 2: In vitro fermentation

Fermentations were performed using a batch technique (Rira et al., 2019). The donor animals were four Texel wethers fitted with a ruminal cannula and weighed on average 80.7 ± 6.9 kg. Wethers were fed daily 900 g hay (natural grassland based on *Dichanthium* spp.) divided into equal amounts at 0700 and 1900 h. Wethers were adapted to the diet for 2 weeks before being used as donors. Four series of 24-h incubations were performed, one per wether. In each series, control *Dichanthium* and the six tannin-rich forages were anaerobically incubated in duplicate as described (Rira et al., 2019).

Gas production was measured at 24 h using a pressure transducer. After recording pressure, a gas sample (5 mL) was taken for methane analysis. For VFA determination, 0.8 mL of filtrate was mixed with 0.5 mL of 4 mg/mL crotonic acid and 20 mg/mL metaphosphoric acid in 0.5 M HCl and frozen at –20°C until analysis.

### Chemical analyses

Feeds and feed residues in bags were analysed according to the Association of Official Analytical Chemists (AOAC, 2005). Organic matter in feeds was determined by ashing at 550 °C for 6 h (AOAC method number 923.03). Nitrogen (N) in feeds and feed residues was determined by the Dumas method (crude protein (CP) = N × 6.25, AOAC method number 992.15). For residues at 24, 48 and 96 h, residues of the 3 cows were pooled to get enough residue for analysis. Cell wall components in feeds (NDF, ADF, and ADL) were determined using sodium sulphite, without heat-stable amylase and including residual ash (AOAC methods number 200.04 and 973.18). Enzymatic dry matter digestibility was estimated in feeds by hydrolysis with pepsin in 0.1 N HCl then with fungal cellulose (Aufrere and Michalet-Doreau, 1988).

Soluble CT, protein-bound CT and fibre-bound CT fractions in tannins-rich plants samples were extracted and analysed according to Terrill et al. (1992) as previously detailed (Rira et al., 2019). The concentration of CT in soluble, protein-bound and fibre-bound fractions was calculated with a standard calibration curve of purified quebracho tannins.

For HT, the rhodanine method was used for determination of gallotannins in feeds (Inoue and Hagerman, 1988). The potassium iodate (KIO3) method was used to estimate total HT (gallotannins and ellagitannins) in feeds (Hartzfeld et al., 2002). Details of these two methods are mentioned in (Rira et al., 2019). Ellagitannins were calculated using the difference between total HT and gallotannins.

After in vitro fermentation, gas composition was determined by gas chromatography (Micro GC 3000A; Agilent Technologies, Les Ulis, France) within 2 h after sampling. Gas molar concentration was calibrated using a certified standard. Volatile fatty acids were analysed by gas chromatography using crotonic acid as internal standard on a Perkin Elmer Clarus 580 GC (Perkin Elmer, Courtaboeuf, France) equipped with a Stabilwax-DA column (30 m by 0.53 mm i.d.) (Morgavi et al., 2013).

### Microbial analysis: DNA extraction and sequencing strategy

DNA was extracted following the protocol described by Yu and Morrison (2004). Total genomic DNA was sent to Roy J. Carver Biotechnology Center (Illinois, USA) for fluidigm amplification and Illumina sequencing using primers targeting bacterial 16S rRNA gene (V3-V5 region), archaeal 16S rRNA gene, fungal ITS2 and 18S rRNA gene for protozoa, as described (Saro et al., 2018, Popova et al., 2019).

#### Microbial bioinformatics analyses

Raw sequencing data were trimmed for quality (Phred score > 25), expected amplicon length (570 nt for bacteria 16S rRNA gene, 457 nt for archaea 16S rRNA gene, 660 nt for 18S rRNA gene and 356 nt for ITS) and maximum 5 primer mismatches. Sequences were analysed using computational pipelines as described (Saro et al., 2018). On average per sample, we obtained 28 045 (±6 457) reads of bacterial 16S rRNA gene, 23 718 (±4 726) for archaeal 16S rRNA gene and 18 645 (±2 696) for eukaryotic 18S rRNA gene. For fungi, most samples had a low number of reads, precluding further comparative analysis between samples. As forward and reverse reads for bacterial 16S rRNA gene amplicons reads and eukaryotic 18S rRNA gene amplicons reads were not overlapping, sequence data were analysed using IM Tornado pipeline (Jeraldo et al., 2014) and taxonomy assigned according to Silva v128. Archaeal sequences were analysed following standard QIIME pipeline (Caporaso et al., 2010) and taxonomy assigned with RIM DB (Seedorf et al., 2014).

OTU tables were analysed in R using the package “vegan” (Oksanen et al., 2016). Diversity indices (Shannon, Simpson, Richness and Evenness) were computed using implemented functions and statistical differences were tested using the non-parametric Kruskal-Wallis test to evaluate the effect of plant at each incubation time. For β-diversity analysis, OTU tables were rarefied to an even depth: 3460 reads for bacteria, 4103 for Archaea and 1000 for protozoa. Bray-Curtis method was used for computing dissimilarity indices with the vegdist function. Principal component analysis (PCoA) was performed on dissimilarity matrices with prcomp function. Permutational multivariate analysis of variance was performed using Adonis function. Correlations were computed using vegan’s function “corr” and “Hmisc” and “corrplot” packages were used for plotting correlation matrices. Differential abundance analysis to evaluate the effect of plant at each incubation time were done with MicrobiomeAnalyst (Chong et al., 2020) using default data filtering (4 minimum count, 20% prevalence, low variance filter: 10% inter-quantile range), relative log expression (RLE) for normalisation and the metagenomeSeq package (Paulson et al., 2013).

### Degradation and fermentation data: calculations and statistical analyses

Rumen degradability (D) of DM and N were calculated using an exponential model with lag time (Denham et al., 1989):
D (t) = a + b (1 – e-c(t-L))
where D is the degradation after t hours of rumen incubation; t = hours of rumen incubation (0, 3, 6, 12, 24, 48 and 96 h); a = rapidly degradable fraction (%); b = slowly degradable fraction (%); c = degradation rate constant of the b fraction (h^-1^); L= lag time before the beginning of degradation of the b fraction (h).

The nonlinear procedure (PROC NLIN) of SAS v9.4 (SAS Inst. Inc., Cary, NC, USA) was used to fit degradation data to the model. This model was chosen after a preliminary comparison of models with or without lag time, because lag time was higher than 3 h for 4 of the 6 feeds, and because the model with lag time globally led to lower sum of squares than the model without lag time. The theoretical degradability in the rumen derived from the model (TDm) was calculated from the equation:
TDm = a + [b × c /(c + kp)]

where kp is the passage rate of solid contents out of the rumen. A unique value of 0.04 h^-1^ was used for all feeds. Rumen degradability of NDF did not fit to the exponential model due to a linear degradation rate for *Calliandra calothyrsus* and to erratic variations for other two forages. For NDF calculations of degradability, we used instead a stepwise method (Kristensen et al., 1982):
TDs = Σ_(i = 0 to n)_ (Dt_(i+1)_ - Dt_i_) × p (t_i_, t_i+1_)

where (Dt_(i+1_)-Dt_i_) is the amount of feed degraded between times t_i+1_ and t_i_, and p (t_i_, t_i+1_) is the proportion of feed remaining in the rumen between times t_i_ and t_i+1_ with pt_i_ = e^−kpti^.

From VFA production in the in vitro experiment, fermented organic matter (FOM) was calculated by the stoichiometric equation of Demeyer and Van Nevel (1975):
FOM = 162 (0.5 acetate + 0.5 propionate + butyrate + valerate) where FOM is expressed in mg and VFA in mmol.

Data of both experiments were submitted to the same mixed model using the MIXED procedure of SAS including feed (n = 6 for in situ experiment and n = 7 for in vitro experiment) as fixed effect and animal (n = 3 for in situ experiment and n = 4 for in vitro experiment) as random effect. Differences between feeds were analysed using the Tukey t-test. Effects were declared significant when P < 0.05. Principal component analyses were performed using Minitab^®^ version 17 software (Minitab Inc., State College, PA). Average values for each of the 6 tannin-rich plants were included for 19 variables: 7 from chemical composition, 5 from experiment 1 and 7 from experiment 2.

## Results

### Forage characteristics

The chemical composition of forages is shown in Table 1. Tannin-containing plants have higher concentration of CP and lower concentration of fibre than the control forage, except *Musa* spp. The enzymatic DM digestibility test showed high values (~70%) for *Acacia nilotica* pods, *Leucaena leucocephala, Gliricidia sepium* and *Manihot esculenta* leaves, whereas digestibility of *Musa* spp. and *Calliandra calothyrsus* leaves was much lower at 40% or less. There were marked differences among forages in both the total amount of CT and the fraction that tannins were associated with (Table 1). For all forages, bound CT were predominantly (≥ 60%) linked to protein compared to fibre. The proportion of free CT was particularly large in *Calliandra calothyrsus* leaves (54% of total CT) and *Acacia nilotica* pods (58% of total CT). In addition, *Acacia nilotica* pods were particularly rich in HT (350 g/kg DM versus less than 33 g/kg DM for the other plants). For the others forages consisting in plant leaves, the HT content was minor. For all forages, ellagitannins were predominant representing ≥ 80% of total HT.

**Table 1.**
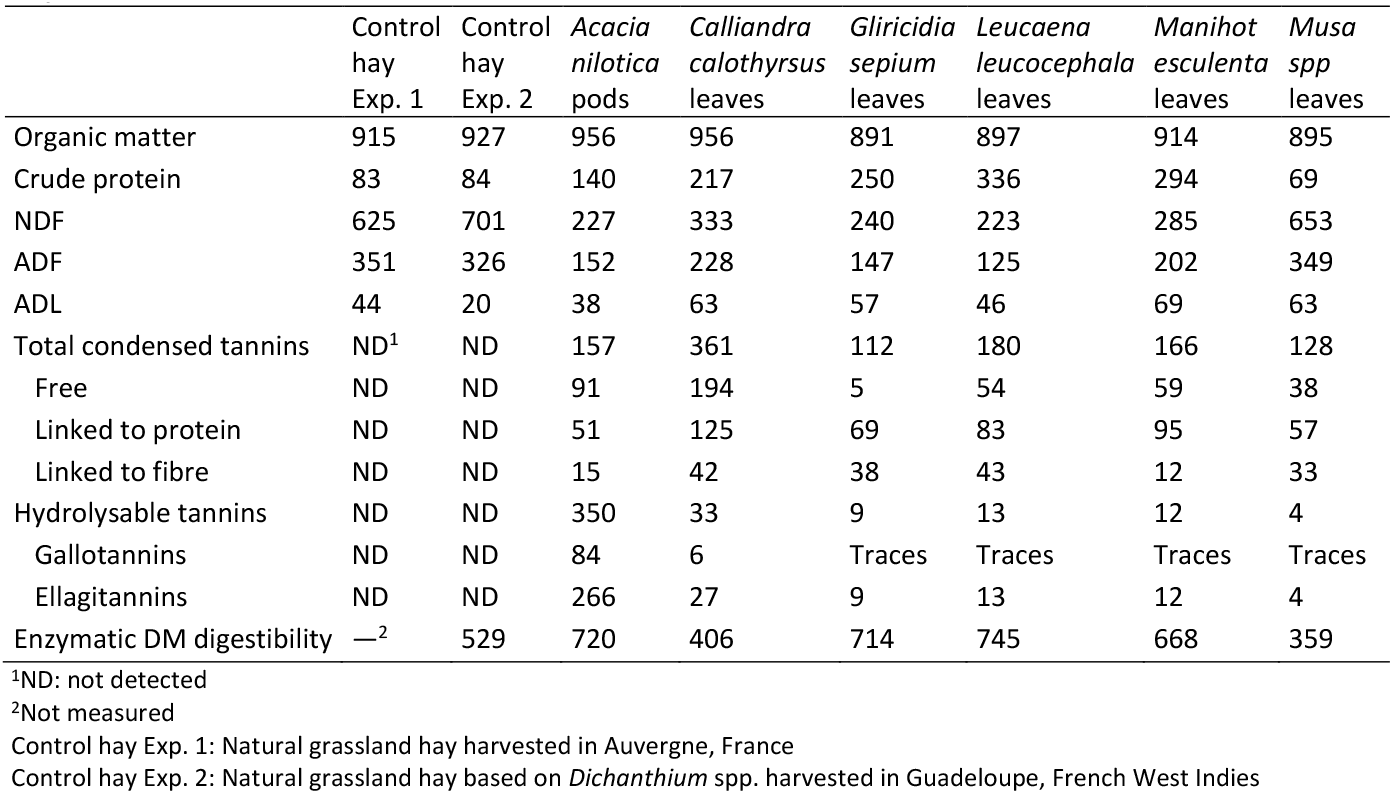
Chemical composition and enzymatic digestibility, in g/kg DM, of plants used in experiments

### In situ dry matter, N and NDF degradability

Dry matter degradability was similar for *Acacia nilotica* pods, *Gliricidia sepium, Manihot esculenta*, and *Leucaena leucocephala* with values around 65%, whereas only a third of *Musa* spp. and *Calliandra calothyrsus* leaves were degraded (Table 2). However, for forages presenting similar degradability, there were differences in the proportion of degraded fractions and rate of degradation. *Acacia nilotica* pods and *Gliricidia sepium* had a higher proportion of soluble fraction (a), whereas *Leucaena leucocephala* and *Manihot esculenta* leaves had a higher proportion of potentially degradable fraction (b). *Musa* spp. and *Calliandra calothyrsus* had lower values for the (a) and (b) fractions than the other forages. Although they have similar degradability, *Musa* spp. leaves had a low proportion of slowly degraded fraction (b) at 14%, whereas *Calliandra calothyrsus* leaves were the most slowly degraded (0.022% h^-1^).

The N degradability showed similar values and ranking than DM degradability. Notwithstanding, there was a higher variation among forages with *Manihot esculenta* leaves presenting the highest overall degradability due to its high proportion of soluble fraction (a) and high rate of degradation (c). For both DM and N, the theoretical degradability calculated according to the model was similar to values obtained by a stepwise calculation. The rumen degradability of NDF was calculated with the stepwise method only because the exponential method with lag time failed to produce suitable models for most forages. Compared to N, NDF was generally less degraded. In most plants around one third of NDF was degraded in the rumen; notable exceptions were *Acacia nilotica* pods with 14% disappearance and *Calliandra calothyrsus* with only 6%.

### Condensed tannins, N and NDF disappearance

Results of 24-h disappearance of tannins, are presented in Table 3, together with N and NDF disappearance to assess if there is a relationship between these parameters. The disappearance of N and NDF at 24 h was calculated from degradability values above to have the same point in time as tannins. After 24 h in the rumen, free CT disappeared completely or almost completely (~98%) in all forages. Among those plants exhibiting a high proportion of protein-bound CT, the disappearance of this fraction was variable with up to 93% loss in *Gliricidia sepium* and no disappearance in *Calliandra calothyrsus*. For this latter forage, it is noted that N disappeared at 24 h was ~32% but it had also the highest amount of CT linked to protein (Table 1). For fibre-bound CT, the average disappearance at 24 h was ~80% and, despite numerical differences, did not differ significantly between forages (P> 0.05). The disappearance of total CT reflected the differences observed in the disappearance of the various fractions and ranged from 58% for *Calliandra calothyrsus* leaves to 95% for *Gliricidia sepium*.

**Table 2.** Dry matter, N and NDF degradability^1^ of tropical tannin-rich plants in the rumen

|  | *Acacia*<br>*nilotica* | *Calliandra*<br>*calothyrsus* | *Gliricidia*<br>*sepium* | *Leucaena*<br>*leucocephala* | *Manihot*<br>*esculenta* | *Musa*<br>*spp* | SEM | P Value |
| --- | --- | --- | --- | --- | --- | --- | --- | --- |
| Dry matter degradability |  |  |  |  |  |  |  |  |
| a (%) | 53.5<sup>a</sup> | 22.1<sup>e</sup> | 49.5<sup>b</sup> | 37.3<sup>c</sup> | 39.2<sup>c</sup> | 27.7<sup>d</sup> | 0.88 | < 0.001 |
| b (%) | 30.4<sup>b</sup> | 25.2<sup>b2</sup> | 27.6<sup>b</sup> | 45.7<sup>a</sup> | 41.4<sup>a</sup> | 14.0<sup>c</sup> | 1.86 | < 0.001 |
| c (h<sup>-1</sup>) | 0.038<sup>ab</sup> | 0.022<sup>b</sup> | 0.170<sup>ab</sup> | 0.075<sup>ab</sup> | 0.189<sup>a</sup> | 0.027<sup>b</sup> | 0.0337 | 0.013 |
| L (h) | 4.24<sup>ab</sup> | 0.00<sup>b</sup> | 5.84<sup>a</sup> | 2.68<sup>ab</sup> | 5.95<sup>a</sup> | 4.90<sup>ab</sup> | 1.595 | 0.026 |
| TDm<sup>3</sup> (%) | 65.2<sup>a</sup> | 30.9<sup>c</sup> | 65.6<sup>a</sup> | 63.3<sup>b</sup> | 66.0<sup>a</sup> | 32.2<sup>c</sup> | 0.46 | < 0.001 |
| TDs<sup>2</sup> (%) | 65.7<sup>a</sup> | 32.0<sup>b</sup> | 65.1<sup>a</sup> | 63.8<sup>a</sup> | 65.5<sup>a</sup> | 33.2<sup>b</sup> | 0.70 | < 0.001 |
| N degradability |  |  |  |  |  |  |  |  |
| a (%) | 40.1<sup>b</sup> | 24.7<sup>e</sup> | 37.8<sup>bc</sup> | 35.6<sup>c</sup> | 47.5<sup>a</sup> | 29.7<sup>d</sup> | 0.74 | < 0.001 |
| b (%) | 58.6<sup>a</sup> | 45.1<sup>ab</sup> | 47.3<sup>a</sup> | 54.6<sup>a</sup> | 45.0<sup>ab</sup> | 13.4<sup>b</sup> | 6.29 | 0.004 |
| c (h<sup>-1</sup>) | 0.031<sup>bc</sup> | 0.013<sup>c</sup> | 0.076<sup>b</sup> | 0.054<sup>bc</sup> | 0.155<sup>a</sup> | 0.025<sup>bc</sup> | 0.0115 | < 0.001 |
| L (h) | 1.88<sup>bc</sup> | 0.00<sup>c</sup> | 5.95<sup>a</sup> | 0.52<sup>c</sup> | 5.40<sup>ab</sup> | 8.54<sup>a</sup> | 0.912 | < 0.001 |
| TDm<sup>2</sup> (%) | 63.7<sup>c</sup> | 33.2<sup>d</sup> | 61.8<sup>c</sup> | 65.7<sup>b</sup> | 76.2<sup>a</sup> | 33.3<sup>d</sup> | 0.57 | < 0.001 |
| TDs<sup>2</sup> (%) | 64.1<sup>c</sup> | 34.7<sup>e</sup> | 61.1<sup>d</sup> | 66.2<sup>b</sup> | 75.7<sup>a</sup> | 34.1<sup>e</sup> | 0.64 | < 0.001 |
| NDF degradability |  |  |  |  |  |  |  |  |
| TDs<sup>2</sup> (%) | 14.5<sup>c</sup> | 6.3<sup>d</sup> | 28.4<sup>b</sup> | 30.0<sup>b</sup> | 35.8<sup>a</sup> | 30.7<sup>ab</sup> | 1.27 | < 0.001 |

<sup>a-d</sup>Values within a row with different superscripts differ significantly at P<0.05, n= 3.
<sup>1</sup> Degradation D was modelled according to the equation $D(t) = a + b(1 - e^{-c(t+L)})$ where a is the rapidly degraded fraction, b the slowly degraded fraction, c the rate of degradation of fraction b, t the time of incubation in the rumen and L the lag time.
<sup>2</sup> n= 2
<sup>3</sup> TDm : theoretical degradability calculated according to the model ; TDs : theoretical degradability calculated according to a stepwise calculation.

**Table 3.** Rumen disappearance (%) of condensed tannins, N and NDF from tropical tannin-rich plants after 24 h of incubation

|  | *Acacia nilotica* | *Calliandra calothyrsus* | *Gliricidia sepium* | *Leucaena leucocephala* | *Manihot esculenta* | *Musa spp* | SEM | P Value |
| --- | --- | --- | --- | --- | --- | --- | --- | --- |
| Condensed tannins | 73.0<sup>d</sup> | 58.2<sup>e</sup> | 94.9<sup>a</sup> | 77.8<sup>cd</sup> | 88.6<sup>ab</sup> | 84.3<sup>bc</sup> | 2.10 | <0.001 |
| Free | 98.3 | 97.4 | 100.0 | 100.0 | 100.0 | 100.0 | - | - |
| Linked to protein | 21.3<sup>d</sup> | -7.8<sup>e</sup> | 92.9<sup>a</sup> | 58.0<sup>c</sup> | 83.0<sup>ab</sup> | 72.7<sup>bc</sup> | 3.24 | <0.001 |
| Linked to fibre | 83.0 | 60.8 | 98.0 | 87.9 | 76.4 | 86.4 | 8.23 | 0.10 |
| N | 65.3<sup>b</sup> | 31.7<sup>c</sup> | 68.8<sup>b</sup> | 73.9<sup>b</sup> | 89.1<sup>a</sup> | 33.8<sup>c</sup> | 2.93 | 0.001 |
| NDF | 11.5<sup>ab</sup> | 5.1<sup>b</sup> | 31.6<sup>ab</sup> | 33.8<sup>ab</sup> | 43.4<sup>a</sup> | 27.8<sup>ab</sup> | 8.21 | 0.012 |

<sup>a-d</sup>Values within a row with different superscripts differ significantly at P<0.05.

### Microbial community attached to tannin-rich plants

We studied ruminal microbial communities attached to plants after 3 and 12 h of incubation in the rumen. Incubation times were chosen in order to catch the biphasic primary and secondary colonisation process described in temperate plants (Elliott et al., 2018). Based on the known differences in communities between these two phases (Mayorga et al., 2016, Elliott et al., 2018), most results are presented by incubation time to better identify the effect of plants.

For anaerobic fungi, most samples had a low number of reads and no downstream analysis was made. However, it is noted that the only plant that was consistently colonised by anaerobic fungi was *Musa* with a threefold increase from 3 to 12 h reaching more than 4000 reads on average (Supplementary Figure 1). For protozoa, *Leucaena leucocephala* had low numbers of reads but it was considered a characteristic of the plant and these samples were included in downstream analysis.

Changes on alpha diversity indices were more marked for bacteria at 3 h (Supplementary Table 1). *Calliandra calothyrsus* had high alpha diversity values that remained numerically higher than for other plants at 12 h. Whereas, archaeal indices differed more at 12 h than at 3 h; *Acacia nilotica* and, to a lesser degree, *Calliandra calothyrsus* were the plants with the lowest values.

Principal coordinate analysis plots showed differences in community structure that were influenced by the type of plant (Figure 1). Permanova analyses highlighted significant differences between plants for all microbial communities at 3 h (Adonis P < 0.001; R^2^= 0.53, 0.43 and 0.59 for bacteria, archaea and protozoa, respectively). These differences remained at 12 h for bacteria (Adonis R^2^= 0.69, P < 0.001) and archaea (Adonis R^2^= 0.48, P < 0.001). At 3 h, the first component separated bacterial communities attached to *Acacia nilotica, Calliandra calothyrsus* and *Leucaena leucocephala* from *Musa, Manihot esculenta*, and the control hay (same hay fed to cows). However, at 12 h, only *Musa* and control hay grouped together clearly separated from the other plants. For archaea, *Acacia nilotica* and *Calliandra calothyrsus* were separated from other plants both at 3 and 12 h of incubation. Whereas, for protozoa, *Leucaena leucocephala* was clearly separated from all other plants at 3 h, undoubtedly due to the low number of reads recovered from this plant; and, at 12 h not clear grouping of plants was observed in agreement with permanova results. The bacterial and archaeal communities correlated with some chemical features of plants; NDF and ADF contents influenced the bacterial community structure and total concentration of tannins, both CT and HT, had a stronger influence on the archaeal community structure (Supplementary Figure 2).

**Figure 1.**
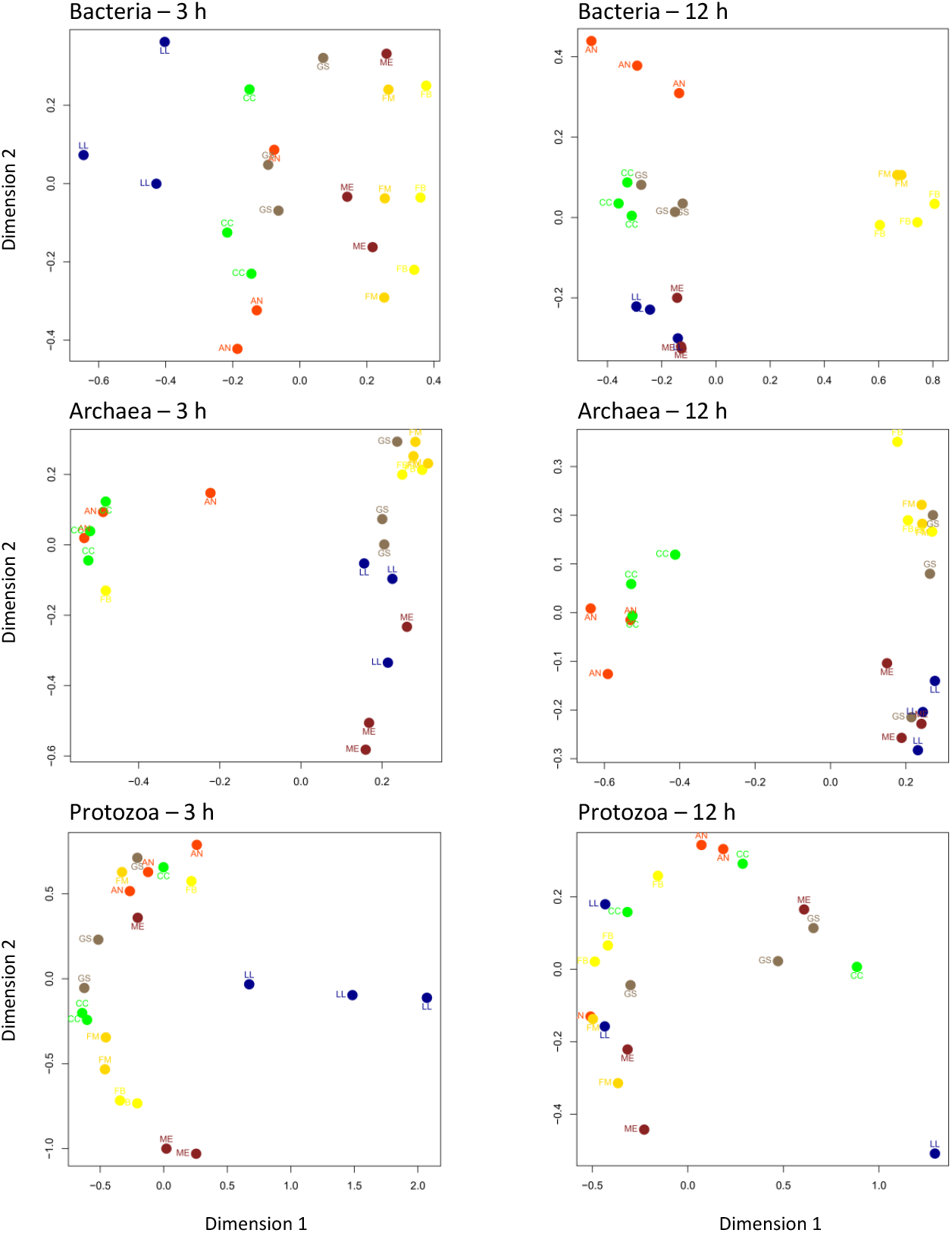
Distribution of bacterial, archaeal and protozoal communities associated to tropical tannin-rich plants after 3 and 12 h of in situ incubation in the rumen (PCoA). 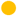 = control hay (no tannin); 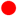 = *Acacia nilotica*; 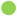 = *Calliandra calothyrsus*; 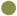 = *Gliricidia sepium*; 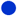 = *Leucaena leucocephala*; 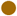 = *Manihot esculenta*; 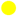 = *Musa spp*.

Differential abundance analyses showed a numerically higher proportion of Proteobacteria for plants richer in tannins at 3 h of incubation (Table 4). At lower taxonomical levels, the family *Rhodospirillaceae* from the Proteobacteria was particularly abundant in *Calliandra calothyrsus*, whereas the γ-proteobacterium *Pantoea* sp. was more abundant in *Calliandra calothyrsus* and *Leucaena leucocephala* (Supplementary Table 2). In contrast, the phylum Fibrobacteres was more abundant in *Musa* and control hay, which were the plants containing a higher amount of fibre and less tannins. The differences were particularly striking at 12 h of incubation with ~25% of sequences belonging to Fibrobacteres in these two plants compared to values as low as 3% for *Acacia nilotica* (Table 4). Results for families and genera levels are shown in Supplementary Tables 2 to 5. *Lachnospiraceae* were proportionally more abundant in *Calliandra calothyrsus, Leucaena leucocephala* and *Gliricidia sepium* at 3 h of incubation and generally more abundant in tannin-rich plants at 12 h of incubation. Representative genera of this family such as *Butyrivibrio* and *Oribacterium* showed the same trend. There were no marked differences in relative abundance of archaea (Supplementary Tables 6 and 7) and, for protozoa, the main change was observed in the relative abundance of *Isotricha* spp. that was higher at 3 h in *Acacia nilotica;* differences were less marked at 12 h. In accord with these results, there was a positive correlation between *Isotricha* spp. and the levels of HT and TCT (Supplementary Tables 8 and 9, and Supplementary Figure 2).

**Table 4.** Relative abundance of rumen bacterial phyla colonizing tropical tannin-rich plants incubated in the rumen of cows (n= 3) for 3 and 12 h

|  | Control<sup>1</sup> | *Acacia nilotica* | *Calliandra calothyrsus* | *Gliricidia sepium* | *Leucaena leucocephala* | *Manihot esculenta* | *Musa spp* | P value | FRD<sup>2</sup> |
| --- | --- | --- | --- | --- | --- | --- | --- | --- | --- |
| After 3 h of incubation |  |  |  |  |  |  |  |  |  |
| *Firmicutes* | 47.61 | 41.16 | 43.47 | 57.90 | 60.08 | 43.58 | 50.26 | 0.554 | 0.727 |
| *Bacteroidetes* | 36.59 | 37.89 | 42.87 | 32.39 | 29.41 | 42.71 | 32.71 | 0.375 | 0.687 |
| *Fibrobacteres* | 11.53 | 2.12 | 3.87 | 5.19 | 3.50 | 8.13 | 12.88 | 0.089 | 0.327 |
| *Spirochaetae* | 1.53 | 1.68 | 3.24 | 1.47 | 2.30 | 3.39 | 1.57 | 0.464 | 0.727 |
| *Actinobacteria* | 1.52 | 6.46 | 2.52 | 1.75 | 1.25 | 0.92 | 1.84 | 0.221 | 0.607 |
| *Proteobacteria* | 0.52 | 9.77 | 3.15 | 0.70 | 2.95 | 0.43 | 0.33 | 0.015 | 0.083 |
| *Tenericutes* | 0.46 | 0.84 | 0.74 | 0.40 | 0.34 | 0.63 | 0.22 | 0.868 | 0.868 |
| *Chloroflexi* | 0.25 | 0.08 | 0.14 | 0.19 | 0.15 | 0.21 | 0.19 | 0.595 | 0.727 |
| After 12 h of incubation |  |  |  |  |  |  |  |  |  |
| *Firmicutes* | 38.54 | 55.98 | 45.32 | 54.81 | 43.48 | 35.15 | 40.42 | 0.592 | 0.609 |
| *Bacteroidetes* | 30.78 | 29.22 | 32.76 | 27.63 | 34.39 | 33.63 | 28.41 | 0.093 | 0.249 |
| *Fibrobacteres* | 26.76a | 2.72d | 5.88c | 11.09b | 13.93b | 11.02b | 23.15a | 0.006 | 0.045 |
| *Spirochaetae* | 1.94 | 5.17 | 7.53 | 3.20 | 5.37 | 13.89 | 7.13 | 0.609 | 0.609 |
| *Tenericutes* | 0.93 | 5.31 | 7.50 | 2.57 | 2.45 | 5.93 | 0.38 | 0.076 | 0.249 |
| *Actinobacteria* | 0.65 | 0.32 | 0.37 | 0.36 | 0.10 | 0.19 | 0.31 | 0.550 | 0.609 |
| *Proteobacteria* | 0.25 | 1.24 | 0.46 | 0.22 | 0.20 | 0.11 | 0.11 | 0.155 | 0.309 |
| *Chloroflexi* | 0.16 | 0.05 | 0.18 | 0.10 | 0.07 | 0.09 | 0.09 | 0.484 | 0.609 |

<sup>1</sup>Control: Natural grassland hay harvested in Auvergne, France
<sup>2</sup>FDR= false discovery rate post-hoc adjustment

Correlation analyses considering the chemical composition of plants, including the various fractions of tannins, and the most abundant (≥ 1%) OTUs only show a few highlighted negative and positive associations. At 3 h, a *Succiniclasticum_uncl*. OTU was negatively correlated to TCT and HT and a *Chistensenellaceae* R7 OTU was positively associated to the fibre-linked fraction of CT. However, at 12 h, the same *Chistensenellaceae* R7 OTU and a *Prevotella* OTU were negatively correlated to the protein linked fraction of CT. A *Fibrobacter* OTU was negatively correlated to HT, whereas *Ruminococcaceae*_NK4A214_group, *Rikenellaceae*_RC9_gut_group_uncl. and a *Butyrivibrio*_2_uncl. were positively correlated to CT (total, free and protein linked) (Figure 2).

**Figure 2.**
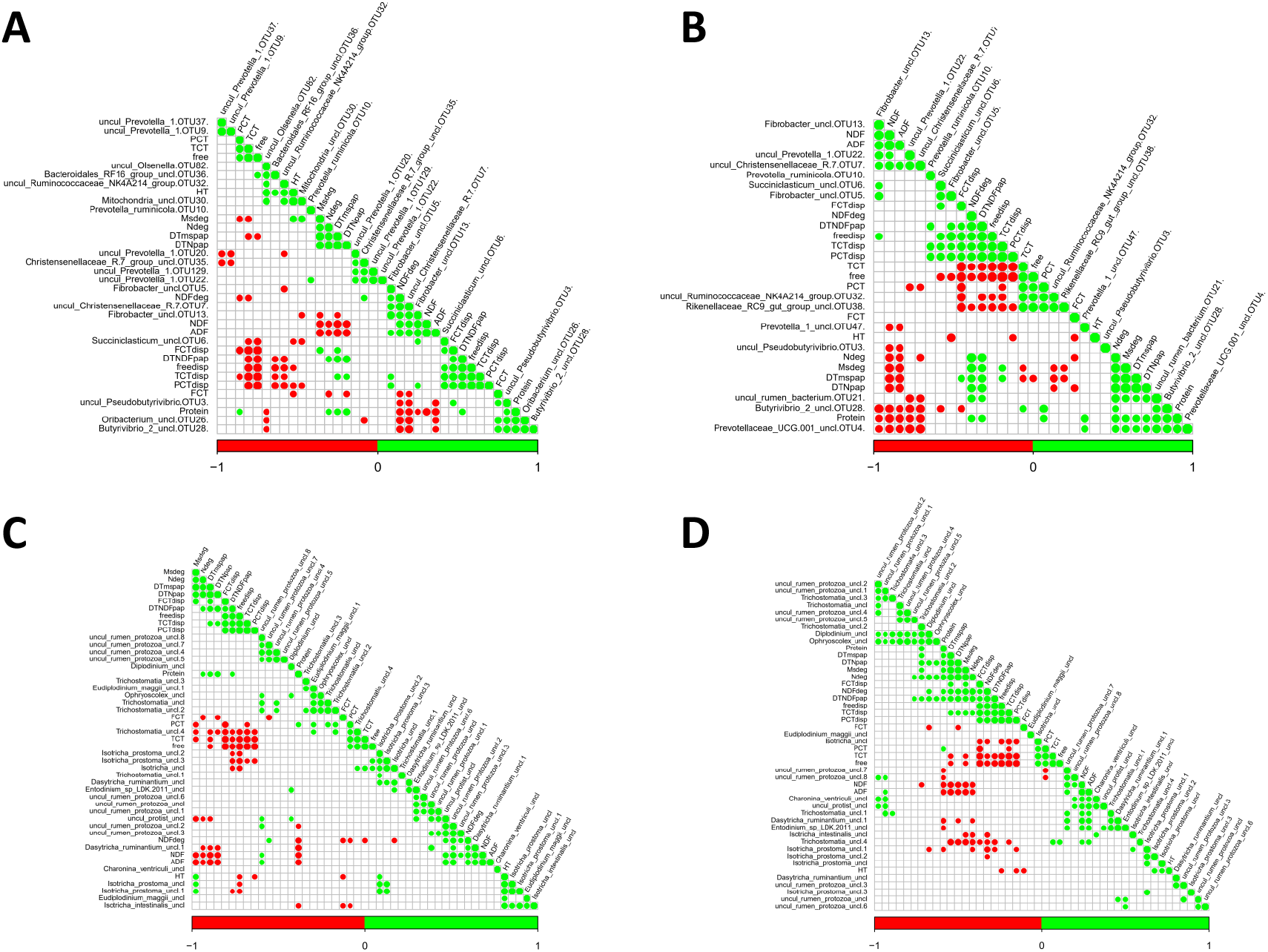
Correlations between microbial species and characteristics of tropical tannin-rich plants after 3 and 12 h of incubation of feeds in the rumen. A: bacteria at 3 h of incubation; B: bacteria, 12 h of incubation; C: protozoa, 3 h of incubation; D: protozoa, 12 h of incubation. For bacteria, operational taxonomic units with abundance > 1% are shown. Green and red dots indicate positive and negative significant correlations (P<0.05), respectively. Dry matter degradability (Msdeg), Nitrogen degradability (Ndeg), Theoretical degradability of dry matter calculated according to a stepwise calculation (TDmspap), Theoretical degradability of nitrogen calculated according to a stepwise calculation (TDNpap), Fibre linked condensed tannins disappearance (FCTdisp), Theoretical degradability of NDF calculated according to a stepwise calculation (TDNDFpap), Free condensed tannins disappearance (freedisp), Total condensed tannins disappearance (TCTdisp), Proteins linked condensed tannins disappearance (PCTdisp), Fibre linked condensed tannins (FCT), Proteins linked condensed tannins (PCT), Total condensed tannins (TCT), Free condensed tannins (free), NDF degradability (NDFdeg), Hydrolysable tannins (HT).

### In vitro rumen fermentation of forages

The in vitro incubation of each tannin-rich plant separately allowed to obtain additional information on their effects on rumen function and methane production. All tannin-rich plants had lower production of gas than control (natural grassland hay based on Dichanthium spp.; Table 5). *Musa* spp. and *Calliandra calothyrsus* were those that produced less gas. Only *Musa* spp., *Calliandra calothyrsus* and *Acacia nilotica* had lower VFA production and FOM than control (P < 0.05). All tannin-rich plants produced more acetate and less butyrate than control, resulting in the absence of difference in the ratio of acetate or acetate+butyrate to propionate between control and tannin-rich plants. Compared to control, methane production when expressed as mL/24 h was reduced for *Leucaena leucocephala*, and to a greater extent for *Musa* spp., *Calliandra calothyrsus* and *Acacia nilotica*. When methane production was expressed per 100 mM of VFA produced, only *Acacia nilotica* and *Musa* spp. decreased production (P < 0.05) compared to control.

**Table 5.**
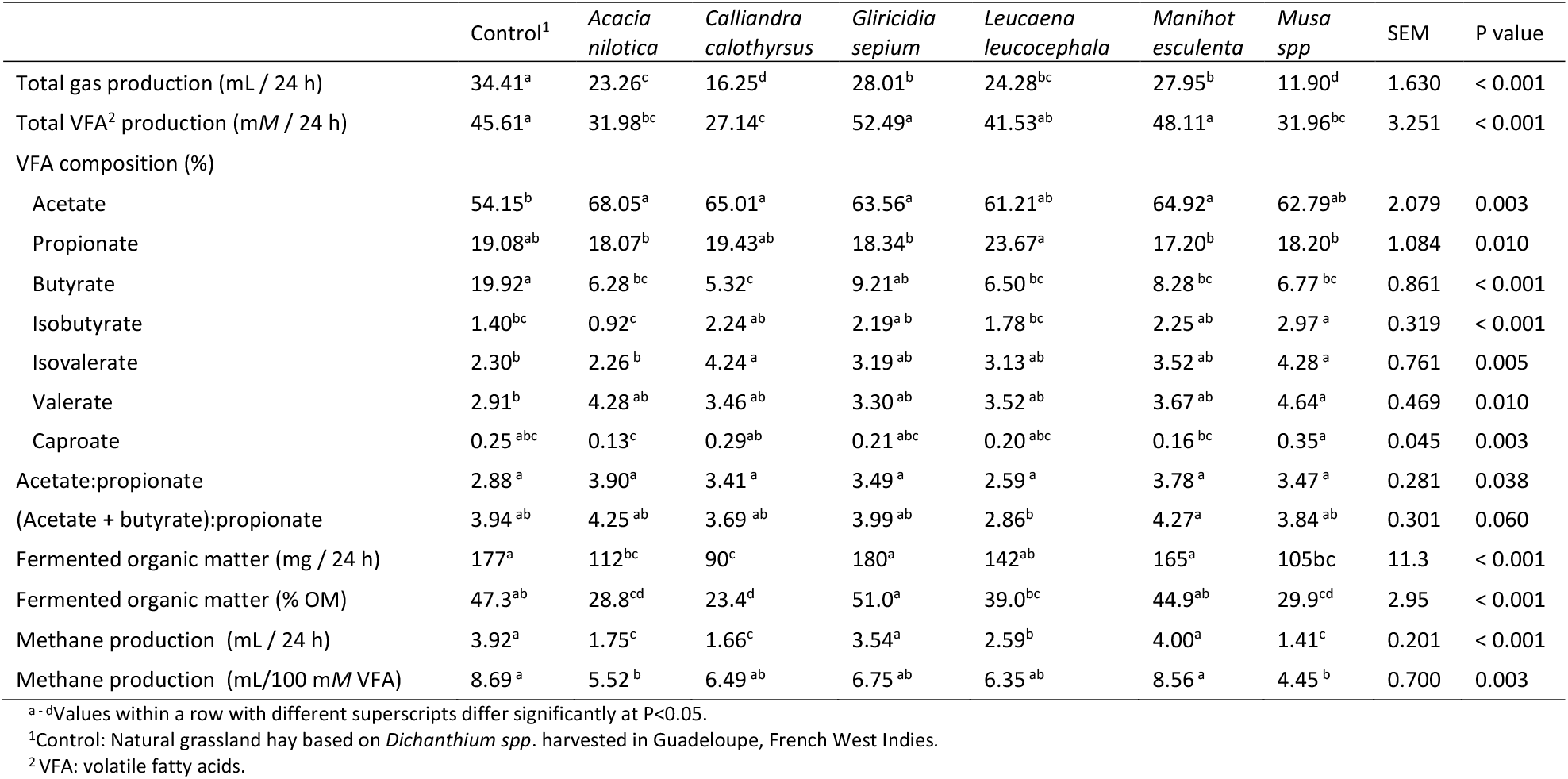
*In vitro* ruminal fermentation of tropical tannin-rich plants

The relationships between the content and type of tannins in plants and the in vitro and in situ parameters were further explored through a principal component analysis (Supplementary Figure 3) that showed that indicators of the extent of ruminal degradation and fermentation, including methane, were opposed on the first axis to total CT content and CT fractions, but not to HT.

## Discussion

All plant species used in this experiment were rich in CT. The amount of CT was within those reported in the literature although mostly on the higher end. The higher CT values in this work can be explained by the analytical method, involving a double-extraction procedure and different solvents for different fractions, potentially yielding a higher amount of extracted tannins than faster methods. Four plants had a higher amount of protein-bound tannins than free tannins whereas two plants (*Acacia nilotica* and *Calliandra calothyrsus*) had a higher amount of free tannins. It is generally reported, even for plants used in this study such as *Leucaena leucocephala* and *Calliandra calothyrsus* that free CT are higher than protein-bound CT that in turn is higher than fibre-bound CT (Terrill et al., 1992, Jackson et al., 1996, Dentinho and Bessa, 2016). However, similar to total CT, there are important divergences in the literature (Perez-Maldonado and Norton, 1996, Rubanza et al., 2005).

In animal nutrition studies, the HT content of plant species is seldom measured and when done it is generally by using non-specific methods, e.g. they are often calculated by the difference between total tannins and CT. We used methods that specifically measured the content of total HT and gallotannins. Among the plants used in our study only *Acacia nilotica* was rich in HT. Goel et al. (2015) also reported high values of HT for this plant (186 g/kg DM, estimated by the difference between total tannins and CT). The other plants used in our study are recognised sources of CT but their HT content is seldom reported. Our results show that these plants rich in CT are also a source of HT, which albeit minor can also have a biological effect.

### Degradation of tannin-rich plants in the rumen - relationship with tannin content

*Calliandra calothyrsus* and *Musa* spp. had a lower DM and N degradability (~33%) than the other plants (~65%). However, there is no a single reason that may explain these differences. *Calliandra calothyrsus* had a markedly low NDF degradability at 6% but it was the disappearance of condensed tannins, representing one third of plant weight, that affected the calculation of DM degradation. Whereas, the low DM degradability of *Musa* spp. is mainly explained by the high proportion of NDF (65% on a DM base). In both cases the rate constant c was low. In contrast, the high degradability of *Leucaena leucocephala, Gliricidia sepium, Acacia nilotica*, and *Manihot esculenta* was due to the high N degradability, to the extensive disappearance of CT, and to the total disappearance of HT from bags. Disproving our hypothesis, we did not observe any relationship either between N degradability and total or protein-bound tannin content, or between NDF degradability and total or fibre-bound tannin content. Similarly, the absence of relationship between total extractible tannins of plants and DM disappearance in situ of seven temperate browses was reported by Khazaal et al. (1993). This contrasts with in vitro degradation results obtained with 72 African browses where protein and NDF degradability were negatively correlated with soluble CT but not with insoluble CT (Rittner and Reed, 1992). The difference with our results may be explained either by the low number of forages in our experiment or by the methodology (in vitro *vs* in situ).

### Rumen disappearance of tannins from plants

Information on tannin disappearance in the digestive tract of ruminants is scarce. Hydrolysable tannins can be degraded by rumen microbes (Brooker et al., 1994, Goel et al., 2005). However, as HT would be washed out of the bags, we did not attempt to quantify them after incubation.

Most authors agree that there is no evidence for free CT degradation by microbes in the rumen (McSweeney et al., 2001b, Patra et al., 2012). Notwithstanding, the complete rumen disappearance of free CT observed is logical because these compounds are water-soluble and are washed out of the bags. A nearly complete (99%) disappearance of free CT of *Calliandra calothyrsus* between mouth and faeces was reported by Perez-Maldonado and Norton (1996).

The disappearance of protein-bound CT varied largely between plants from 21 to 98%. This variation may be due to differences in strength of binding between proteins and tannins (Le Bourvellec and Renard, 2019). As stated above, no relationship was observed between disappearance of proteins and protein-bound CT except for *Calliandra calothyrsus* that had a low N degradation and no disappearance of protein-bound tannins. For this plant, a negative value for disappearance was even obtained. This may be due to a technical problem as interactions with the insoluble matrix, proteins, polysaccharides, and other plant polymers can decrease the solubility of tannins in the extractant, resulting in an underestimation of tannin content in feeds (Dentinho and Bessa, 2016). Another possible reason is a linkage of dietary free CT with proteins. According to Hagerman (1989), if CT are present in excess, all proteins available are bound to tannins, leading to insoluble complexes. This is the case for *Calliandra calothyrsus* which contains more tannins than proteins (361 vs 217 g/kg, respectively).

The fibre-bound CT represented a small proportion of the total CT content of the forages used in our study. Their disappearance in the rumen varied between 61 and 98% but differences between forages were not significant.

### Tannin-rich plants and methane production

Tannin concentration is often considered a critical factor affecting ruminal fermentation (Patra and Saxena, 2011). Correspondingly, we generally observed that the CT content of plants was negatively associated to in situ rumen degradability and to in vitro fermentation parameters including methane production. However, the effect on methane of some plants could also be due to parameters other than CT. As protein concentration affects the volume of gas produced by the bicarbonate buffer in the in vitro fermentation system, we will discuss methane production normalised by VFA for assessing the impact of tannin-rich plants independently of the extent of gas production. Compared to control, *Acacia nilotica* and *Musa* spp. reduced methane production but probably not for the same reasons. For *Musa* spp. the effect cannot be ascribed to the amount of tannins as this plant has a low concentration compared to others. The effect could be due to other secondary compounds such as polyphenols that are present in *Musa* spp. leaves in large amounts (Marie-Magdeleine et al., 2010). For *Acacia nilotica*, tannins are the most plausible cause of methane inhibition as this plant contains CT but is especially rich in HT. We previously showed that HT could be more efficient than CT for reducing methane (Rira et al., 2019). Other authors have also reported that HT can reduce methane production without compromising overall rumen fermentation, but to date only extracts were studied (Bhatta et al., 2009, Hassanat and Benchaar, 2013, Jayanegara et al., 2015). It is likely that HT do not interfere with rumen fermentation because they do not bind to protein or fibre and do not have an inhibitory effect on microbes that can also degrade HT in the rumen (Patra et al., 2012).

In this study, *Gliricidia sepium* and *Manihot esculenta* did not decrease methane production. In previous studies, *Gliricidia sepium* was less effective than *Leucaena leucocephala* and *Manihot esculenta* for decreasing methane *in vitro* (Rira et al., 2015) and *in vivo* (Archimede et al., 2016). The absence of effect of *Manihot esculenta* on methane production in this study is unexpected, but is consistent with the large variability of response of methane production to CT for a same plant (Piluzza et al., 2014).

### Tannin-rich plants modulate adherent rumen microbes

The colonising microbial community differed between plants and was influenced by tannins as well as other chemical features such as fibre content. As plants were incubated in the same rumen environment and exposed to the same microbiota there is no doubt that the plant itself is selecting for their adherent microbiota, at least in the initial colonisation stages monitored in this work. The process is probably a combination of microbial tolerance to tannins’ toxicity (Frutos et al., 2004) and substrate preferences. Our results bring new insight on how methane production in the rumen is affected by tannin-rich plants and concurs with the reported absence of a relationship between chemical structure and biological effect of tannins including methane production (McAllister et al., 2005, Naumann et al., 2018).

There is no equivalent published information on the attachment of rumen microbes to these plants for a straight comparison. In contrast, the effect of tannin-rich plants or extracts added to the diet have been reported in several studies.

The fungal community was targeted in our study but it was not further analysed as a low number of reads was recovered. Nevertheless, *Musa* spp. with one of the lowest amounts of tannins among the tested plants was the only one colonised by fungi. This result could be interpreted as an inhibitory effect of tannins on fungi as reported previously (Muhammed et al., 1995). For protozoa, it is noted that *Isotricha* spp. were more abundant on *Acacia nilotica* samples at 3 h of incubation suggesting that this genus was attracted to HT particularly abundant in this plant. This result is only comparable within this study as the pore size of the in sacco incubation bags may obstruct the free passage of protozoa but it can help explain the variable effect of different tannin sources on protozoa reported in the literature (Patra and Saxena, 2011).

The plants’ characteristics clearly influenced the structure of the attached bacterial community. Initially, each plant seems to harbour a distinct community of primary colonisers. Then, at 12 h when most of soluble tannins are no longer interfering, the communities in tannin-rich plants became more similar but distinctly separated from communities found in the control forage without tannin and the low-tannin containing *Musa*. For archaea the pattern is different with only *Acacia nilotica* and *Calliandra calothyrsus* clearly separating from other plants. This indicate that the plant effect is more important on colonising bacteria than on archaea, which is logical as the latter rely on metabolic end products from other microbes. The different archaeal structure for tannin-rich *Acacia nilotica* and *Calliandra calothyrsus* could reflect a toxicity threshold attained at the biofilm microenvironment level that needs to be proved.

Our results bring new evidence indicating that some of the differences reported on rumen microbes when tannins are supplemented to the diet are due to changes in the colonisation and development of feed-attached communities. The low proportion of *Fibrobacter* attached to tannin-rich plants may explain why this fibrolytic bacterium is often affected in supplemented animals (McSweeney et al., 2001a, Diaz Carrasco et al., 2017, Harun et al., 2017, Salami et al., 2018). On the other hand, some bacteria that were positively associated to plants rich in tannins could provide new avenues of exploration for improving the nutritional value of these forages through modulation of the microbiota.

## Conclusion

We used an integrated approach for assessing the effect of tannin-rich plants on rumen processes. Tannin-rich plants have contrasting proportions of tannins bound to protein or fibre. However, there was no relationship between the amount and disappearance of bound tannins and the rumen degradability of protein or fibre of the plants studied in this work, except for *Calliandra calothyrsus* that was extremely rich in CT. Tannins present in plants restrained the colonization of certain rumen microbial populations known to play a role in the degradation of recalcitrant feeds. These results expand our understanding of the effects of tannins in the rumen opening the way to improve their use in the diet of ruminants.

## Data, script and code availability

Sequencing data are available in the Sequence Read Archive (SRA) under accession ID PRJNA554299. Scripts and other data can be found at https://doi.org/10.15454/8J2O5C

## Acknowledgements

The authors thank the staff of UE 1414 Herbipôle for their technical collaboration, animal care and management, A. Torrent and L. Genestoux for their contribution in the experimental setup and sample analysis, and A. Roche (intern student at INRAE) for her contribution in the analysis of condensed tannins. We are grateful to the INRAE MIGALE bioinformatics platform (http://migale.jouy.inra.fr) for providing computational resources and A. Bernard for initial bioinformatic analysis.

Version 3 of this preprint has been peer-reviewed and recommended by Peer Community In Animal Science (https://doi.org/10.24072/pci.animsci.100009).

## Appendix

**Supplementary Table 1.**
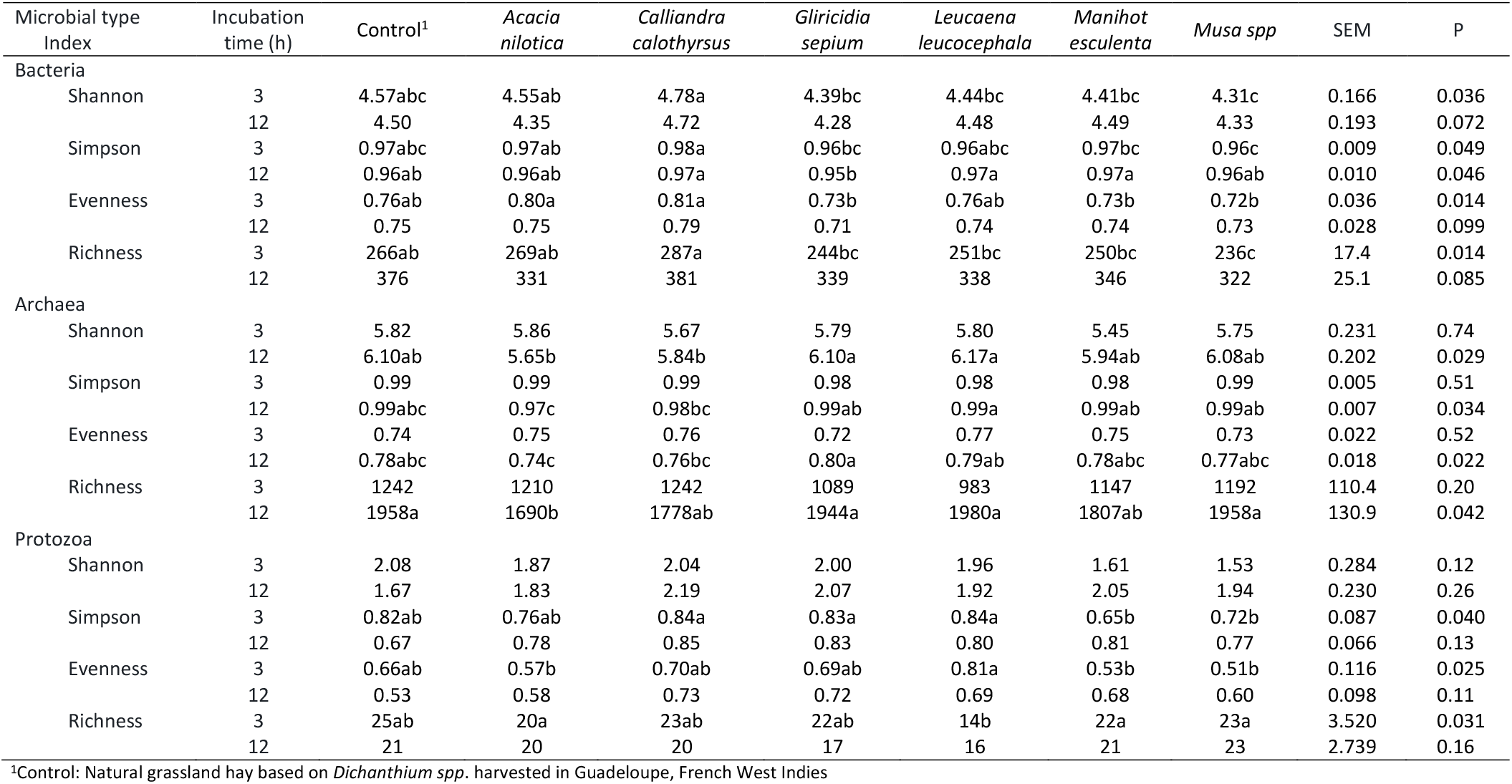
Diversity indices for bacterial, archaeal and protozoal populations colonising tropical tannin-rich plants incubated in the rumen of cows (n= 3) for 3 and 12 h, statistical analysis was performed using the non parametric Kruskal-Wallis test

**Supplementary Table 2.** Proportional abundance of bacterial families colonising tropical tannin-rich plants incubated in the rumen of cows (n= 3) for 3 h

|  | Control<sup>1</sup> | *Acacia*<br>*nilotica* | *Calliandra*<br>*calothyrsus* | *Gliricidia*<br>*sepium* | *Leucaena*<br>*leucocephala* | *Manihot*<br>*esculenta* | *Musa spp* | P value<sup>2</sup> | FRD |
| --- | --- | --- | --- | --- | --- | --- | --- | --- | --- |
| *Prevotellaceae* | 28.9 | 24.2 | 30.6 | 26.2 | 23.0 | 37.7 | 27.5 | 0.042 | 0.147 |
| *Lachnospiraceae* | 15.6 | 12.0 | 21.2 | 30.2 | 41.8 | 16.9 | 14.0 | 0.010 | 0.045 |
| *Christensenellaceae* | 15.3 | 15.9 | 10.9 | 14.1 | 7.5 | 15.6 | 20.2 | 0.635 | 0.846 |
| *Fibrobacteraceae* | 11.5 | 2.1 | 3.9 | 5.2 | 3.5 | 8.1 | 12.9 | 0.005 | 0.030 |
| *Ruminococcaceae* | 7.3 | 8.0 | 6.0 | 5.2 | 3.8 | 5.9 | 9.4 | 0.911 | 0.966 |
| *Streptococcaceae* | 4.1 | 0.2 | 0.1 | 0.8 | 0.3 | 0.1 | 0.5 | 0.916 | 0.966 |
| *Rikenellaceae* | 4.0 | 4.9 | 4.7 | 2.2 | 2.5 | 2.0 | 2.3 | 0.626 | 0.846 |
| *Acidaminococcaceae* | 3.5 | 1.5 | 2.5 | 6.4 | 2.3 | 3.7 | 5.3 | 0.042 | 0.147 |
| *Bacteroidales_BS11_gut_group* | 1.7 | 3.1 | 2.1 | 1.4 | 1.5 | 1.0 | 1.4 | 0.762 | 0.927 |
| *Spirochaetaceae* | 1.6 | 1.8 | 3.3 | 1.5 | 2.4 | 3.4 | 1.6 | 0.017 | 0.069 |
| *Coriobacteriaceae* | 1.5 | 6.5 | 2.5 | 1.7 | 1.2 | 0.9 | 1.8 | 0.165 | 0.421 |
| *Bacteroidales_RF16_group* | 1.3 | 4.5 | 4.4 | 2.0 | 1.7 | 1.1 | 1.0 | 0.444 | 0.654 |
| *Veillonellaceae* | 0.7 | 2.3 | 2.1 | 0.7 | 3.7 | 0.4 | 0.4 | 0.309 | 0.633 |
| *Bacteroidales_S24_7_group* | 0.6 | 0.5 | 0.4 | 0.4 | 0.3 | 0.5 | 0.3 | 0.369 | 0.633 |
| *Clostridiales_vadinBB60_group* | 0.5 | 0.2 | 0.1 | 0.0 | 0.1 | 0.1 | 0.1 | 0.074 | 0.207 |
| Family_XIII | 0.4 | 0.5 | 0.5 | 0.4 | 0.3 | 0.2 | 0.2 | 0.758 | 0.927 |
| *Rhodospirillaceae* | 0.3 | 0.8 | 2.3 | 0.4 | 0.3 | 0.2 | 0.2 | 0.004 | 0.030 |
| *Anaeroplasmataceae* | 0.3 | 0.6 | 0.6 | 0.3 | 0.3 | 0.5 | 0.2 | 0.407 | 0.633 |
| *Anaerolineaceae* | 0.2 | 0.1 | 0.1 | 0.2 | 0.1 | 0.2 | 0.2 | 0.917 | 0.966 |
| *Bacteroidales_UCG_001* | 0.2 | 0.9 | 0.3 | 0.1 | 0.3 | 0.1 | 0.2 | 0.287 | 0.633 |
| *Succinivibrionaceae* | 0.2 | 0.6 | 0.4 | 0.3 | 0.9 | 0.2 | 0.1 | 0.392 | 0.633 |
| *Erysipelotrichaceae* | 0.1 | 0.3 | 0.2 | 0.0 | 0.1 | 0.0 | 0.1 | 0.370 | 0.633 |
| *Bacteroidetes_unclassified* | 0.1 | 0.1 | 0.2 | 0.2 | 0.2 | 0.3 | 0.1 | 0.966 | 0.966 |
| *Bacteroidales_unclassified* | 0.0 | 0.0 | 0.0 | 0.0 | 0.1 | 0.0 | 0.1 | 0.916 | 0.966 |
| *Mollicutes_unclassified* | 0.0 | 0.1 | 0.0 | 0.0 | 0.0 | 0.0 | 0.0 | 0.392 | 0.633 |

<sup>1</sup>Control: Natural grassland hay based on *Dichanthium spp.* harvested in Guadeloupe, French West Indies
<sup>2</sup>Differential abundance was estimated with the metagenomeSeq package (Paulson et al., 2013)

**Supplementary Table 3.** Proportional abundance of bacterial families colonising tropical tannin-rich plants incubated in the rumen of cows (n= 3) for 12 h

|  | Control<sup>1</sup> | *Acacia nilotica* | *Calliandra calothyrsus* | *Gliricidia sepium* | *Leucaena leucocephala* | *Manihot esculenta* | *Musa spp* | P value<sup>2</sup> | FRD |
| --- | --- | --- | --- | --- | --- | --- | --- | --- | --- |
| *Fibrobacteraceae* | 26.9 | 2.7 | 5.9 | 11.1 | 13.9 | 11.0 | 23.1 | 0.022 | 0.099 |
| *Prevotellaceae* | 20.8 | 24.5 | 21.5 | 21.8 | 28.0 | 27.8 | 24.7 | 0.571 | 0.727 |
| *Christensenellaceae* | 12.7 | 9.9 | 7.3 | 7.7 | 7.3 | 6.3 | 12.9 | 0.112 | 0.253 |
| *Lachnospiraceae* | 12.5 | 39.6 | 30.7 | 39.9 | 29.8 | 22.3 | 14.5 | 0.309 | 0.504 |
| *Ruminococcaceae* | 8.2 | 2.7 | 4.9 | 3.3 | 3.1 | 4.6 | 9.7 | 0.004 | 0.021 |
| *Rikenellaceae* | 7.7 | 2.4 | 7.0 | 2.5 | 2.7 | 2.8 | 2.6 | 0.002 | 0.015 |
| *Acidaminococcaceae* | 2.4 | 0.9 | 1.1 | 3.2 | 2.1 | 1.5 | 2.5 | 0.670 | 0.787 |
| *Spirochaetaceae* | 1.9 | 5.1 | 7.0 | 3.2 | 5.3 | 13.9 | 7.2 | 0.940 | 0.976 |
| *Clostridiales_vadinBB60_group* | 1.4 | 0.1 | 0.9 | 0.2 | 0.4 | 0.1 | 0.2 | <0.001 | <0.001 |
| *Bacteroidales_S24_7_group* | 1.2 | 1.7 | 3.1 | 2.2 | 2.5 | 2.0 | 0.6 | 0.044 | 0.133 |
| *Anaeroplasmataceae* | 0.7 | 3.2 | 4.8 | 2.0 | 1.9 | 4.2 | 0.2 | 0.069 | 0.169 |
| *Coriobacteriaceae* | 0.6 | 0.3 | 0.4 | 0.4 | 0.1 | 0.2 | 0.3 | 0.420 | 0.629 |
| *Bacteroidales_BS11_gut_group* | 0.6 | 0.3 | 0.7 | 0.5 | 0.2 | 0.3 | 0.3 | 0.027 | 0.102 |
| *Veillonellaceae* | 0.5 | 1.3 | 0.3 | 0.3 | 0.5 | 0.2 | 0.2 | 0.030 | 0.102 |
| Family_XIII | 0.4 | 0.2 | 0.3 | 0.3 | 0.1 | 0.1 | 0.2 | 0.708 | 0.796 |
| *Bacteroidales_RF16_group* | 0.3 | 0.3 | 0.4 | 0.6 | 0.5 | 0.2 | 0.2 | 0.521 | 0.727 |
| *Streptococcaceae* | 0.3 | 1.2 | 0.0 | 0.2 | 0.0 | 0.0 | 0.1 | 0.000 | 0.000 |
| *Bacteroidales_UCG_001* | 0.2 | 0.1 | 0.1 | 0.0 | 0.0 | 0.0 | 0.2 | 0.989 | 0.989 |
| *Rhodospirillaceae* | 0.2 | 0.1 | 0.3 | 0.1 | 0.1 | 0.1 | 0.1 | 0.317 | 0.504 |
| *Anaerolineaceae* | 0.2 | 0.0 | 0.2 | 0.1 | 0.1 | 0.1 | 0.1 | 0.845 | 0.912 |
| *Succinivibrionaceae* | 0.1 | 0.8 | 0.1 | 0.1 | 0.1 | 0.1 | 0.0 | <0.001 | <0.001 |
| *Bacteroidetes_unclassified* | 0.1 | 0.2 | 0.4 | 0.1 | 0.7 | 0.7 | 0.1 | 0.153 | 0.317 |
| *Erysipelotrichaceae* | 0.1 | 0.0 | 0.1 | 0.0 | 0.0 | 0.0 | 0.0 | 0.547 | 0.727 |
| *Mollicutes_unclassified* | 0.1 | 2.0 | 2.5 | 0.2 | 0.3 | 1.4 | 0.0 | 0.592 | 0.727 |
| *Bacteroidales_unclassified* | 0.0 | 0.0 | 0.0 | 0.0 | 0.1 | 0.1 | 0.1 | 0.237 | 0.456 |

<sup>1</sup>Control hay: Natural grassland hay based on *Dichanthium spp.* harvested in Guadeloupe, French West Indies
<sup>2</sup>Differential abundance was estimated with the metagenomeSeq package (Paulson et al., 2013)

**Supplementary Table 4.** Proportional abundance of bacterial taxa colonising tropical tannin-rich plants incubated in the rumen of cows (n= 3) for 3 h

|  | Control<sup>1</sup> | *Acacia nilotica* | *Calliandra calothyrsus* | *Gliricidia sepium* | *Leucaena leucocephala* | *Manihot esculenta* | *Musa spp* | P value<sup>2</sup> | FRD |
| --- | --- | --- | --- | --- | --- | --- | --- | --- | --- |
| Uncultured_rumen_bacterium | 62.8 | 60.6 | 60.7 | 67.3 | 57.7 | 67.6 | 67.0 | 0.498 | 0.755 |
| *Fibrobacter*_unclassified | 9.8 | 1.7 | 3.0 | 4.8 | 3.0 | 7.2 | 10.8 | 0.030 | 0.248 |
| *Streptococcus*_unclassified | 4.2 | 0.2 | 0.1 | 0.8 | 0.3 | 0.1 | 0.5 | 0.451 | 0.747 |
| *Prevotella*_1_unclassified | 4.0 | 3.2 | 4.3 | 3.8 | 2.6 | 4.2 | 3.3 | 0.318 | 0.722 |
| Uncultured_rumen_bacterium_unclassified | 3.0 | 6.6 | 5.4 | 3.3 | 2.6 | 2.5 | 2.2 | 0.878 | 0.934 |
| Uncultured_bacterium | 2.9 | 3.3 | 4.3 | 3.1 | 2.9 | 3.7 | 2.7 | 0.193 | 0.653 |
| *Butyrivibrio*_2_unclassified | 1.8 | 1.3 | 4.5 | 4.4 | 7.0 | 3.3 | 1.4 | 0.003 | 0.041 |
| *Christensenellaceae*_R_7_group_unclassified | 1.4 | 1.7 | 1.5 | 1.5 | 0.9 | 1.6 | 1.5 | 0.622 | 0.864 |
| *Ruminococcus flavefaciens* | 0.8 | 0.8 | 0.5 | 0.5 | 0.4 | 0.9 | 1.6 | 0.705 | 0.904 |
| *Prevotella_ruminicola* | 0.8 | 1.2 | 1.7 | 1.0 | 1.4 | 1.5 | 0.9 | 0.306 | 0.722 |
| *Oribacterium*_unclassified | 0.6 | 0.3 | 1.2 | 1.6 | 3.9 | 0.8 | 0.4 | 0.005 | 0.047 |
| Uncultured_rumen_bacterium_5C0d_11 | 0.5 | 0.2 | 0.3 | 0.2 | 0.3 | 0.3 | 0.6 | 0.694 | 0.904 |
| Uncultured_bacterium_unclassified | 0.5 | 1.9 | 1.3 | 0.4 | 0.7 | 0.3 | 0.5 | 0.821 | 0.933 |
| Unidentified_rumen_bacterium_RC2 | 0.5 | 0.3 | 0.5 | 0.2 | 0.4 | 0.2 | 0.3 | 0.108 | 0.542 |
| Uncultured_rumen_bacterium_4C28d_15_unclassified | 0.4 | 0.2 | 0.1 | 0.0 | 0.1 | 0.1 | 0.1 | 0.389 | 0.747 |
| *Selenomonas*_1_unclassified | 0.4 | 0.5 | 0.8 | 0.4 | 2.0 | 0.2 | 0.1 | 0.921 | 0.940 |
| *Ruminococcus*_sp_HUN007 | 0.4 | 0.3 | 0.3 | 0.1 | 0.0 | 0.1 | 0.8 | 0.663 | 0.895 |
| uncultured_*Lachnospiraceae*_bacterium | 0.4 | 0.2 | 0.3 | 0.5 | 0.4 | 0.6 | 0.7 | 0.139 | 0.612 |
| Bacterium_AC2043 | 0.3 | 0.4 | 0.3 | 0.3 | 0.2 | 0.3 | 0.5 | 0.814 | 0.933 |
| *Bacteroidales*_BS11_gut_group_unclassified | 0.3 | 0.4 | 0.4 | 0.1 | 0.3 | 0.1 | 0.3 | 0.595 | 0.864 |
| Uncultured_rumen_bacterium_3C0d_9 | 0.3 | 0.1 | 0.3 | 0.1 | 0.1 | 0.1 | 0.2 | 0.227 | 0.709 |
| Uncultured_*Prevotellaceae*_bacterium | 0.3 | 0.1 | 0.4 | 0.4 | 0.3 | 0.4 | 0.3 | 0.095 | 0.530 |
| Uncultured_*Clostridium*_sp | 0.3 | 0.1 | 0.1 | 0.1 | 0.1 | 0.2 | 0.5 | 0.294 | 0.722 |
| *Selenomonas_ruminantium*_AB3002 | 0.2 | 1.6 | 1.1 | 0.2 | 1.3 | 0.2 | 0.2 | 0.278 | 0.722 |
| *Butyrivibrio_fibrisolvens* | 0.2 | 0.3 | 0.5 | 0.9 | 1.7 | 0.4 | 0.2 | 0.155 | 0.612 |
| *Lachnospiraceae*_bacterium_AC2012 | 0.2 | 0.2 | 0.1 | 0.4 | 0.4 | 0.1 | 0.1 | 0.361 | 0.747 |
| *Prevotella*_sp_CA17 | 0.2 | 0.2 | 0.3 | 0.3 | 0.2 | 0.3 | 0.1 | 0.943 | 0.943 |
| X_*Eubacterium_ventriosum*_group_unclassified | 0.2 | 0.3 | 0.2 | 0.0 | 0.1 | 0.1 | 0.1 | 0.772 | 0.929 |
| *Lachnospiraceae*_unclassified | 0.2 | 0.1 | 0.1 | 0.1 | 0.2 | 0.2 | 0.1 | 0.919 | 0.940 |
| *Succinivibrio*_unclassified | 0.2 | 0.6 | 0.4 | 0.3 | 0.9 | 0.2 | 0.1 | 0.404 | 0.747 |
| *Lachnospiraceae*_bacterium_NK4A144 | 0.2 | 0.2 | 0.4 | 0.3 | 0.8 | 0.2 | 0.1 | 0.159 | 0.612 |
| Unidentified_rumen_bacterium_RC17 | 0.2 | 0.2 | 0.1 | 0.1 | 0.1 | 0.1 | 0.2 | 0.383 | 0.747 |
| *Rikenellaceae*_RC9_gut_group_unclassified | 0.2 | 0.5 | 0.2 | 0.1 | 0.1 | 0.1 | 0.1 | 0.463 | 0.747 |
| *Lachnospiraceae*_NK3A20_group_unclassified | 0.2 | 0.1 | 0.2 | 0.2 | 0.0 | 0.1 | 0.1 | 0.196 | 0.653 |
| Bacterium_XPD3003 | 0.1 | 0.1 | 0.1 | 0.1 | 0.0 | 0.1 | 0.2 | 0.444 | 0.747 |
| Uncultured_rumen_bacterium_3C0d_20 | 0.1 | 0.1 | 0.0 | 0.2 | 0.1 | 0.2 | 0.2 | 0.460 | 0.747 |
| Unidentified_rumen_bacterium_RFN89 | 0.1 | 0.1 | 0.0 | 0.3 | 0.2 | 0.2 | 0.1 | 0.242 | 0.712 |
| *Roseburia_unclassified* | 0.1 | 0.1 | 0.1 | 0.3 | 0.3 | 0.2 | 0.1 | 0.463 | 0.747 |
| *Bacteroidales_RF16_group_unclassified* | 0.1 | 0.2 | 0.3 | 0.1 | 0.1 | 0.1 | 0.1 | 0.723 | 0.904 |
| *Bacteroidetes_unclassified* | 0.1 | 0.1 | 0.1 | 0.2 | 0.2 | 0.3 | 0.1 | 0.872 | 0.934 |
| *Bacterium_XPB1013* | 0.1 | 0.0 | 0.2 | 0.2 | 0.2 | 0.1 | 0.1 | 0.072 | 0.516 |
| *X_Eubacterium_hallii_group_unclassified* | 0.1 | 0.1 | 0.1 | 0.1 | 0.1 | 0.1 | 0.0 | 0.610 | 0.864 |
| *Ruminococcus_albus* | 0.1 | 0.0 | 0.0 | 0.1 | 0.0 | 0.1 | 0.2 | 0.781 | 0.929 |
| *Bacterium_XPB4001* | 0.1 | 0.3 | 0.3 | 0.1 | 0.1 | 0.1 | 0.0 | 0.306 | 0.722 |
| *Unidentified_rumen_bacterium_RF31* | 0.1 | 0.2 | 0.2 | 0.3 | 0.2 | 0.2 | 0.1 | 0.857 | 0.934 |
| *Unidentified_rumen_bacterium_RFN13* | 0.1 | 0.0 | 0.0 | 0.2 | 0.2 | 0.2 | 0.1 | 0.497 | 0.755 |
| *Bacteria_unclassified* | 0.0 | 0.0 | 0.8 | 0.1 | 0.1 | 0.1 | 0.0 | <0.001 | 0.001 |
| *Clostridiales_vadinBB60_group_unclassified* | 0.0 | 0.5 | 0.1 | 0.0 | 0.2 | 0.0 | 0.0 | 0.090 | 0.530 |
| *Pantoea_unclassified* | 0.0 | 0.1 | 1.1 | 0.0 | 3.0 | 0.0 | 0.0 | 0.001 | 0.018 |

<sup>1</sup>Control: Natural grassland hay based on *Dichanthium spp.* harvested in Guadeloupe, French West Indies
<sup>2</sup>Differential abundance was estimated with the metagenomeSeq package (Paulson et al., 2013)

**Supplementary Table 5.** Proportional abundance of bacterial taxa colonising tropical tannin-rich plants incubated in the rumen of cows (n= 3) for 12 h

|  | Control<sup>1</sup> | *Acacia nilotica* | *Calliandra calothyrsus* | *Gliricidia sepium* | *Leucaena leucocephala* | *Manihot esculenta* | *Musa spp* | P value<sup>2</sup> | FRD |
| --- | --- | --- | --- | --- | --- | --- | --- | --- | --- |
| Uncultured_rumen_bacterium | 53.1 | 52.5 | 54.2 | 56.7 | 47.6 | 51.3 | 55.4 | 0.938 | 0.979 |
| *Fibrobacter*_unclassified | 23.7 | 2.0 | 4.6 | 9.9 | 12.2 | 9.6 | 19.9 | 0.030 | 0.142 |
| Uncultured_bacterium | 2.5 | 6.4 | 3.8 | 2.2 | 3.8 | 5.8 | 2.4 | 0.157 | 0.430 |
| Uncultured_rumen_bacterium_unclassified | 2.0 | 1.9 | 3.3 | 3.1 | 2.7 | 2.2 | 1.1 | 0.202 | 0.482 |
| *Prevotella*_1_unclassified | 2.0 | 3.8 | 4.5 | 3.2 | 6.3 | 3.0 | 2.9 | 0.545 | 0.799 |
| Uncultured_rumen_bacterium_4C28d_15_unclassified | 1.4 | 0.1 | 0.8 | 0.2 | 0.4 | 0.1 | 0.2 | <0.001 | <0.001 |
| *Prevotella_ruminicola* | 1.4 | 0.7 | 1.0 | 2.9 | 1.1 | 2.3 | 1.1 | 0.517 | 0.795 |
| Unidentified_rumen_bacterium_RC2 | 1.3 | 0.5 | 1.0 | 0.3 | 0.3 | 0.3 | 0.5 | 0.131 | 0.394 |
| Uncultured_rumen_bacterium_5C0d_11 | 1.1 | 0.1 | 0.1 | 0.1 | 0.2 | 0.5 | 2.6 | 0.977 | 0.979 |
| *Ruminococcus flavefaciens* | 1.0 | 0.1 | 0.2 | 0.3 | 0.3 | 0.5 | 1.7 | 0.206 | 0.482 |
| Uncultured_Prevotellaceae_bacterium | 0.9 | 2.9 | 3.1 | 2.6 | 5.2 | 4.5 | 2.4 | 0.130 | 0.394 |
| *Christensenellaceae*_R_7_group_unclassified | 0.8 | 1.3 | 0.7 | 0.6 | 0.6 | 0.6 | 0.7 | 0.077 | 0.271 |
| *Ruminococcus*_sp_HUN007 | 0.8 | 0.1 | 0.1 | 0.0 | 0.0 | 0.1 | 1.8 | 0.961 | 0.979 |
| *Butyrivibrio*_2_unclassified | 0.8 | 4.8 | 7.0 | 4.0 | 5.9 | 6.9 | 1.4 | 0.147 | 0.422 |
| Uncultured_Lachnospiraceae_bacterium | 0.6 | 0.4 | 0.5 | 2.5 | 0.9 | 0.5 | 0.7 | 0.757 | 0.924 |
| Uncultured_Clostridium_sp | 0.6 | 0.1 | 0.1 | 0.1 | 0.0 | 0.1 | 0.4 | 0.481 | 0.757 |
| *Oribacterium*_unclassified | 0.5 | 0.4 | 0.4 | 2.0 | 2.4 | 0.4 | 0.3 | 0.778 | 0.924 |
| *Treponema*_2_unclassified | 0.4 | 1.6 | 3.2 | 0.6 | 1.2 | 1.4 | 0.5 | 0.024 | 0.137 |
| Uncultured_rumen_bacterium_3C0d_9 | 0.3 | 0.3 | 0.3 | 0.1 | 0.1 | 0.0 | 0.2 | 0.979 | 0.979 |
| *Prevotella*_sp_CA17 | 0.3 | 0.1 | 0.1 | 0.2 | 0.1 | 0.2 | 0.2 | 0.269 | 0.527 |
| *Streptococcus*_unclassified | 0.3 | 1.1 | 0.0 | 0.2 | 0.0 | 0.0 | 0.1 | <0.001 | <0.001 |
| *Selenomonas ruminantium*_AB3002 | 0.2 | 0.5 | 0.1 | 0.2 | 0.2 | 0.0 | 0.1 | 0.010 | 0.073 |
| bacterium_AC2043 | 0.2 | 0.3 | 0.2 | 0.2 | 0.2 | 0.1 | 0.2 | 0.270 | 0.527 |
| *Butyrivibrio fibrisolvens* | 0.2 | 4.4 | 1.2 | 1.5 | 1.1 | 0.8 | 0.2 | 0.005 | 0.042 |
| Bacterium_XPD3003 | 0.2 | 0.1 | 0.1 | 0.1 | 0.1 | 0.1 | 0.2 | 0.955 | 0.979 |
| *Lachnospiraceae*_unclassified | 0.2 | 0.2 | 0.4 | 0.3 | 0.3 | 0.3 | 0.1 | 0.071 | 0.263 |
| *Lachnospiraceae*_bacterium_AC2012 | 0.2 | 1.4 | 0.3 | 0.8 | 0.2 | 0.1 | 0.2 | <0.001 | 0.004 |
| Uncultured_bacterium_unclassified | 0.2 | 0.0 | 0.3 | 0.1 | 0.1 | 0.1 | 0.1 | 0.032 | 0.142 |
| Unidentified_rumen_bacterium_RFN66 | 0.2 | 0.1 | 0.0 | 0.3 | 0.2 | 0.0 | 0.1 | 0.004 | 0.037 |
| *Selenomonas*_1_unclassified | 0.2 | 0.7 | 0.1 | 0.1 | 0.2 | 0.1 | 0.1 | 0.046 | 0.181 |
| Unidentified_rumen_bacterium_RC11 | 0.2 | 0.2 | 0.2 | 0.1 | 0.1 | 0.1 | 0.2 | 0.735 | 0.924 |
| *X_Eubacterium coprostanoligenes*_group_unclassified | 0.2 | 0.1 | 0.2 | 0.1 | 0.0 | 0.1 | 0.1 | 0.848 | 0.953 |
| *Ruminococcus albus* | 0.1 | 0.1 | 0.1 | 0.0 | 0.1 | 0.1 | 0.3 | 0.694 | 0.911 |
| Unidentified_rumen_bacterium_RFN89 | 0.1 | 0.1 | 0.1 | 0.2 | 0.1 | 0.0 | 0.0 | 0.617 | 0.845 |
| Unidentified_rumen_bacterium_RFN13 | 0.1 | 0.1 | 0.1 | 0.1 | 0.2 | 0.2 | 0.0 | 0.042 | 0.176 |
| *X_Eubacterium hallii*_group_unclassified | 0.1 | 0.2 | 0.1 | 0.1 | 0.1 | 0.0 | 0.1 | 0.085 | 0.282 |
| *Bacteroidales*_BS11_gut_group_unclassified | 0.1 | 0.1 | 0.1 | 0.0 | 0.0 | 0.0 | 0.1 | 0.820 | 0.939 |
| Human_gut_metagenome | 0.1 | 0.1 | 0.2 | 0.0 | 0.1 | 0.1 | 0.1 | 0.643 | 0.861 |
| Uncultured_prokaryote | 0.1 | 0.0 | 0.0 | 0.0 | 0.0 | 0.0 | 0.0 | 0.440 | 0.710 |
| *Succinivibrio*_unclassified | 0.1 | 0.8 | 0.1 | 0.1 | 0.1 | 0.1 | 0.0 | 0.000 | 0.001 |
| *Roseburia*_unclassified | 0.1 | 0.8 | 0.2 | 1.1 | 0.4 | 0.5 | 0.1 | 0.002 | 0.019 |
| Bacterium_XPB1013 | 0.1 | 0.3 | 0.4 | 0.2 | 0.8 | 0.5 | 0.3 | 0.293 | 0.527 |
| *Lachnospiraceae*_bacterium_NK4A144 | 0.1 | 0.7 | 0.3 | 0.3 | 0.5 | 0.1 | 0.1 | 0.023 | 0.137 |
| *Ruminococcus*_1_unclassified | 0.1 | 0.0 | 0.0 | 0.0 | 0.1 | 0.1 | 0.0 | 0.164 | 0.430 |
| *Bacteroidales*_S24_7_group_unclassified | 0.1 | 0.2 | 0.4 | 0.3 | 0.3 | 0.2 | 0.0 | 0.792 | 0.924 |
| Uncultured_rumen_bacterium_3C0d_20 | 0.1 | 0.1 | 0.1 | 0.1 | 0.1 | 0.1 | 0.1 | 0.615 | 0.845 |
| Unidentified_rumen_bacterium_RF31 | 0.1 | 0.5 | 0.2 | 0.4 | 0.2 | 0.2 | 0.1 | 0.002 | 0.019 |
| *X_Eubacterium_ventriosum*_group_unclassified | 0.1 | 0.1 | 0.1 | 0.0 | 0.0 | 0.1 | 0.1 | 0.900 | 0.979 |
| *Lachnospiraceae*_NK3A20_group_unclassified | 0.1 | 0.1 | 0.2 | 0.1 | 0.0 | 0.1 | 0.1 | 0.349 | 0.611 |
| *X_Clostridium_aminophilum* | 0.1 | 0.0 | 0.0 | 0.1 | 0.2 | 0.1 | 0.0 | 0.272 | 0.527 |
| *Bacteroidetes*_unclassified | 0.1 | 0.2 | 0.4 | 0.1 | 0.7 | 0.7 | 0.1 | 0.176 | 0.444 |
| *Mollicutes*_unclassified | 0.1 | 1.8 | 2.4 | 0.2 | 0.3 | 1.4 | 0.0 | 0.289 | 0.527 |
| *Saccharofermentans*_unclassified | 0.0 | 0.0 | 0.1 | 0.0 | 0.0 | 0.0 | 0.1 | 0.361 | 0.615 |
| *Anaeroplasma*_unclassified | 0.0 | 1.5 | 1.7 | 0.3 | 0.2 | 1.4 | 0.0 | 0.279 | 0.527 |
| *Bacteroidales*_unclassified | 0.0 | 0.0 | 0.0 | 0.0 | 0.1 | 0.1 | 0.1 | 0.289 | 0.527 |
| Unidentified_rumen_bacterium_RC17 | 0.0 | 0.1 | 0.1 | 0.1 | 0.1 | 0.1 | 0.2 | 0.951 | 0.979 |
| *X_Eubacterium_rectale*_group_unclassified | 0.0 | 0.0 | 0.1 | 0.1 | 0.0 | 0.2 | 0.1 | 0.760 | 0.924 |
| *Treponema*_sp_S | 0.0 | 0.2 | 0.2 | 0.1 | 0.2 | 0.6 | 0.1 | 0.541 | 0.799 |
| Unidentified | 0.0 | 3.0 | 0.3 | 0.1 | 0.7 | 0.3 | 0.0 | 0.249 | 0.527 |
| *X_Eubacterium_cellulosolvens*_6 | 0.0 | 0.0 | 0.0 | 0.0 | 0.0 | 0.1 | 0.2 | 0.605 | 0.845 |
| Bacterium_ND2006 | 0.0 | 0.2 | 0.1 | 0.1 | 0.2 | 0.0 | 0.0 | 0.026 | 0.138 |
| *Porphyromonadaceae*_unclassified | 0.0 | 0.0 | 0.0 | 0.0 | 0.0 | 0.1 | 0.0 | 0.429 | 0.710 |
| *Prevotella_bryantii*_B14 | 0.0 | 0.1 | 0.1 | 0.3 | 0.1 | 0.3 | 0.0 | 0.779 | 0.924 |

<sup>1</sup>Control: Natural grassland hay based on *Dichanthium* spp. harvested in Guadeloupe, French West Indies
<sup>2</sup>Differential abundance was estimated with the metagenomeSeq package (Paulson et al., 2013)

**Supplementary Table 6.** Proportional abundance of archaea colonising tropical tannin-rich plants incubated in the rumen of cows (n= 3) for 3 h

|  | Control<sup>1</sup> | *Acacia nilotica* | *Calliandra calothyrsus* | *Gliricidia sepium* | *Leucaena leucocephala* | *Manihot esculenta* | *Musa spp* | P value<sup>2</sup> | FRD |
| --- | --- | --- | --- | --- | --- | --- | --- | --- | --- |
| *Methanobrevibacter_gottschalkii*_clade | 61.4 | 65.1 | 63.0 | 65.3 | 66.6 | 65.5 | 69.6 | 0.929 | 0.929 |
| Not_Assigned | 17.5 | 14.3 | 13.9 | 17.9 | 15.1 | 12.9 | 11.2 | 0.772 | 0.836 |
| Group9_sp | 9.2 | 8.7 | 10.8 | 8.8 | 5.3 | 7.7 | 8.6 | 0.406 | 0.627 |
| *Methanobrevibacter_ruminantium*_clade | 6.2 | 4.6 | 6.0 | 4.1 | 4.8 | 5.3 | 5.9 | 0.434 | 0.627 |
| Group10_sp | 2.8 | 1.4 | 2.7 | 1.4 | 7.1 | 5.5 | 1.9 | 0.054 | 0.213 |
| *Methanosphaera_sp_ISO3_F5* | 2.0 | 4.1 | 1.9 | 1.4 | 1.0 | 1.8 | 1.6 | 0.065 | 0.213 |
| *Methanosphaera_stadtmanae* | 0.3 | 0.3 | 0.4 | 0.2 | 0.0 | 0.2 | 0.7 | 0.554 | 0.655 |
| *Methanobrevibacter_oralis* | 0.2 | 0.4 | 0.3 | 0.1 | 0.1 | 0.2 | 0.2 | 0.334 | 0.621 |
| *Methanosphaera_cuniculi* | 0.2 | 0.4 | 0.1 | 0.1 | 0.0 | 0.1 | 0.2 | 0.008 | 0.098 |
| *Methanobrevibacter_smithii* | 0.1 | 0.0 | 0.1 | 0.1 | 0.0 | 0.1 | 0.1 | 0.331 | 0.621 |
| *Methanomicrobium_mobile* | 0.1 | 0.5 | 0.7 | 0.3 | 0.0 | 0.1 | 0.0 | 0.232 | 0.604 |
| *Methanobacterium_alkaliphilum* | 0.1 | 0.0 | 0.0 | 0.0 | 0.0 | 0.1 | 0.0 | 0.038 | 0.213 |

<sup>1</sup>Control: Natural grassland hay based on *Dichanthium spp.* harvested in Guadeloupe, French West Indies<sup>2</sup>Differential abundance was estimated with the metagenomeSeq package (Paulson et al., 2013)**Supplementary Table 7.** Proportional abundance of archaea colonising tropical tannin-rich plants incubated in the rumen of cows (n= 3) for 12 h

**Supplementary Table 7.**
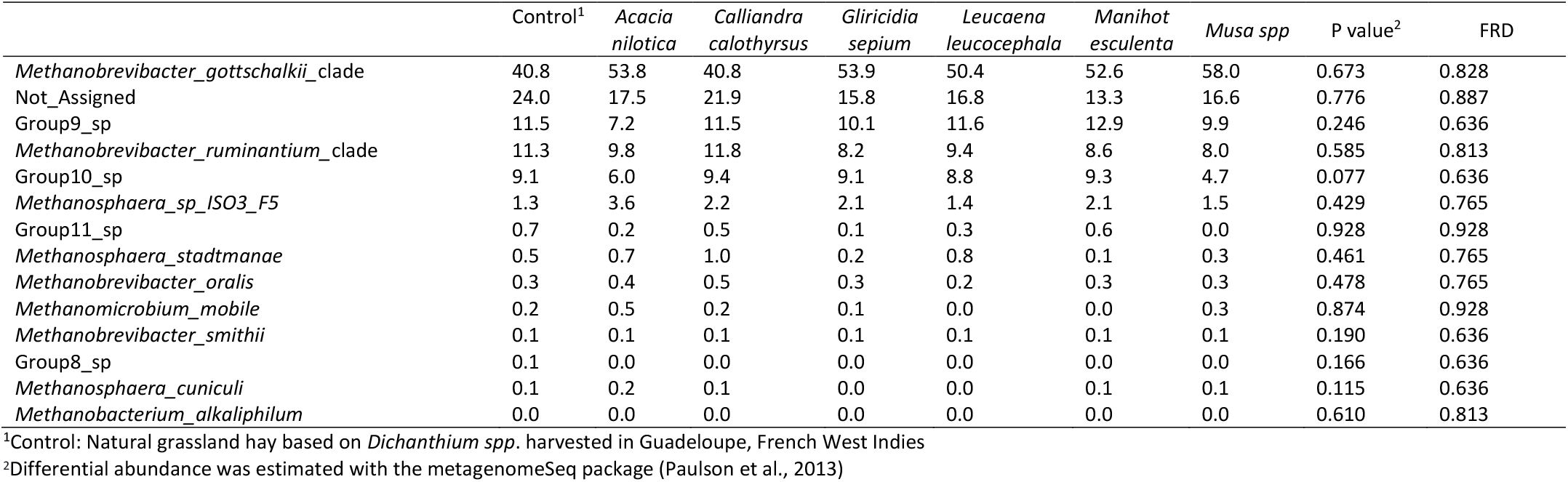
Proportional abundance of archaea colonising tropical tannin-rich plants incubated in the rumen of cows (n= 3) for 12 h

|  | Control<sup>1</sup> | *Acacia nilotica* | *Calliandra calothyrsus* | *Gliricidia sepium* | *Leucaena leucocephala* | *Manihot esculenta* | *Musa spp* | P value<sup>2</sup> | FRD |
| --- | --- | --- | --- | --- | --- | --- | --- | --- | --- |
| *Methanobrevibacter_gottschalkii*_clade | 40.8 | 53.8 | 40.8 | 53.9 | 50.4 | 52.6 | 58.0 | 0.673 | 0.828 |
| Not_Assigned | 24.0 | 17.5 | 21.9 | 15.8 | 16.8 | 13.3 | 16.6 | 0.776 | 0.887 |
| Group9_sp | 11.5 | 7.2 | 11.5 | 10.1 | 11.6 | 12.9 | 9.9 | 0.246 | 0.636 |
| *Methanobrevibacter_ruminantium*_clade | 11.3 | 9.8 | 11.8 | 8.2 | 9.4 | 8.6 | 8.0 | 0.585 | 0.813 |
| Group10_sp | 9.1 | 6.0 | 9.4 | 9.1 | 8.8 | 9.3 | 4.7 | 0.077 | 0.636 |
| *Methanosphaera_sp_ISO3_F5* | 1.3 | 3.6 | 2.2 | 2.1 | 1.4 | 2.1 | 1.5 | 0.429 | 0.765 |
| Group11_sp | 0.7 | 0.2 | 0.5 | 0.1 | 0.3 | 0.6 | 0.0 | 0.928 | 0.928 |
| *Methanosphaera_stadtmanae* | 0.5 | 0.7 | 1.0 | 0.2 | 0.8 | 0.1 | 0.3 | 0.461 | 0.765 |
| *Methanobrevibacter_oralis* | 0.3 | 0.4 | 0.5 | 0.3 | 0.2 | 0.3 | 0.3 | 0.478 | 0.765 |
| *Methanomicrobium_mobile* | 0.2 | 0.5 | 0.2 | 0.1 | 0.0 | 0.0 | 0.3 | 0.874 | 0.928 |
| *Methanobrevibacter_smithii* | 0.1 | 0.1 | 0.1 | 0.1 | 0.1 | 0.1 | 0.1 | 0.190 | 0.636 |
| Group8_sp | 0.1 | 0.0 | 0.0 | 0.0 | 0.0 | 0.0 | 0.0 | 0.166 | 0.636 |
| *Methanosphaera_cuniculi* | 0.1 | 0.2 | 0.1 | 0.0 | 0.0 | 0.1 | 0.1 | 0.115 | 0.636 |
| *Methanobacterium_alkaliphilum* | 0.0 | 0.0 | 0.0 | 0.0 | 0.0 | 0.0 | 0.0 | 0.610 | 0.813 |

<sup>1</sup>Control: Natural grassland hay based on *Dichanthium spp.* harvested in Guadeloupe, French West Indies<sup>2</sup>Differential abundance was estimated with the metagenomeSeq package (Paulson et al., 2013)

**Supplementary Table 8.** Proportional abundance of protozoa associated to tropical tannin-rich plants incubated in the rumen of cows (n= 3) for 3 h

|  | Control<sup>1</sup> | *Acacia nilotica* | *Calliandra calothyrsus* | *Gliricidia sepium* | *Leucaena leucocephala* | *Manihot esculenta* | *Musa spp* | P value | FRD |
| --- | --- | --- | --- | --- | --- | --- | --- | --- | --- |
| Uncultured_rumen_protozoa_unclassified | 63.8 | 28.7 | 51.2 | 51.9 | 60.2 | 79.8 | 79.6 | 0.029 | 0.113 |
| *Isotricha prostoma*_unclassified | 20.8 | 63.1 | 30.2 | 32.6 | 18.3 | 7.5 | 13.6 | <0.001 | 0.001 |
| *Diplodinium*_unclassified | 3.1 | 0.2 | 1.1 | 1.5 | 3.0 | 2.7 | 0.3 | 0.388 | 0.657 |
| *Trichostomatia*_unclassified | 3.1 | 0.7 | 2.9 | 2.1 | 2.7 | 2.4 | 0.8 | 0.802 | 0.935 |
| Uncultured_protist_unclassified | 2.4 | 0.5 | 1.6 | 1.2 | 1.2 | 1.9 | 2.8 | 0.813 | 0.935 |
| *Eudiplodinium maggii*_unclassified | 2.4 | 1.0 | 4.0 | 3.2 | 6.0 | 1.9 | 0.2 | 0.870 | 0.953 |
| *Ophryoscolex*_unclassified | 1.5 | 0.2 | 1.2 | 1.4 | 1.7 | 1.4 | 0.3 | 0.376 | 0.657 |
| *Isotricha intestinalis*_unclassified | 1.3 | 3.3 | 1.0 | 1.1 | 1.3 | 1.0 | 2.0 | <0.001 | <0.001 |
| *Dasytricha ruminantium*_unclassified | 0.9 | 0.5 | 1.2 | 1.5 | 0.2 | 0.6 | 0.3 | 0.206 | 0.475 |
| *Isotricha*_unclassified | 0.7 | 2.0 | 5.6 | 3.3 | 5.5 | 0.7 | 0.2 | 0.095 | 0.273 |

<sup>1</sup>Control: Natural grassland hay based on *Dichanthium spp.* harvested in Guadeloupe, French West Indies
<sup>2</sup>Differential abundance was estimated with the metagenomeSeq package (Paulson et al., 2013)

**Supplementary Table 9.** Proportional abundance of protozoa associated to tropical tannin-rich plants incubated in the rumen of cows (n= 3) for 12 h

|  | Control<sup>1</sup> | *Acacia nilotica* | *Calliandra calothyrsus* | *Gliricidia sepium* | *Leucaena leucocephala* | *Manihot esculenta* | *Musa spp* | P value | FRD |
| --- | --- | --- | --- | --- | --- | --- | --- | --- | --- |
| Uncultured_rumen_protozoa_unclassified | 62.9 | 53.4 | 36.2 | 36.2 | 55.8 | 65.7 | 66.6 | 0.005 | 0.046 |
| *Eudiplodinium maggii*_unclassified | 17.2 | 10.9 | 14.6 | 14.6 | 16.9 | 8.2 | 3.8 | 0.787 | 0.871 |
| *Isotricha prostoma*_unclassified | 10.6 | 29.3 | 33.1 | 33.1 | 19.3 | 10.2 | 21.9 | 0.046 | 0.208 |
| *Diplodinium*_unclassified | 3.3 | 0.3 | 1.2 | 1.2 | 0.4 | 4.2 | 0.4 | 0.663 | 0.871 |
| *Trichostomatia*_unclassified | 3.0 | 1.3 | 2.4 | 2.4 | 2.2 | 4.4 | 1.0 | 0.401 | 0.871 |
| *Isotricha intestinalis*_unclassified | 1.1 | 2.2 | 3.5 | 3.5 | 1.0 | 1.2 | 2.5 | 0.588 | 0.871 |
| Uncultured_protist_unclassified | 0.8 | 0.8 | 1.2 | 1.2 | 0.0 | 3.3 | 3.5 | 0.848 | 0.871 |
| *Ophryoscolex*_unclassified | 0.6 | 0.2 | 0.2 | 0.2 | 0.2 | 0.9 | 0.1 | 0.871 | 0.871 |
| *Isotricha*_unclassified | 0.5 | 1.5 | 7.3 | 7.3 | 3.7 | 1.3 | 0.3 | 0.273 | 0.819 |

<sup>1</sup>Control: Natural grassland hay based on *Dichanthium spp.* harvested in Guadeloupe, French West Indies
<sup>2</sup>Differential abundance was estimated with the metagenomeSeq package (Paulson et al., 2013)

**Supplementary Fig. 1.**
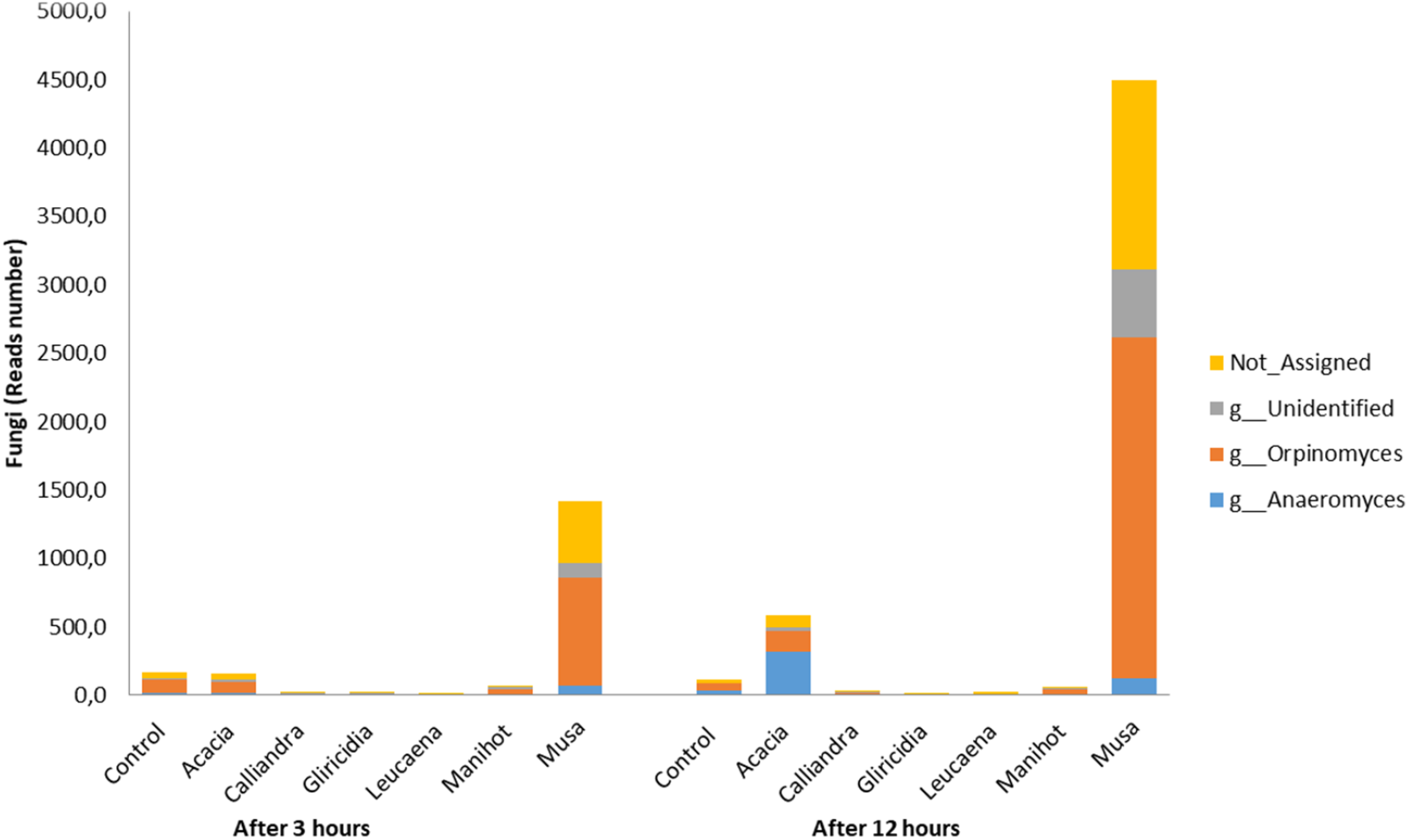
Abundance of anaerobic fungal taxa (ITS amplicon) colonising tropical tannin-rich plants incubated in the rumen of cows (n= 3) for 3 and 12 h

**Supplementary Figure 2.**
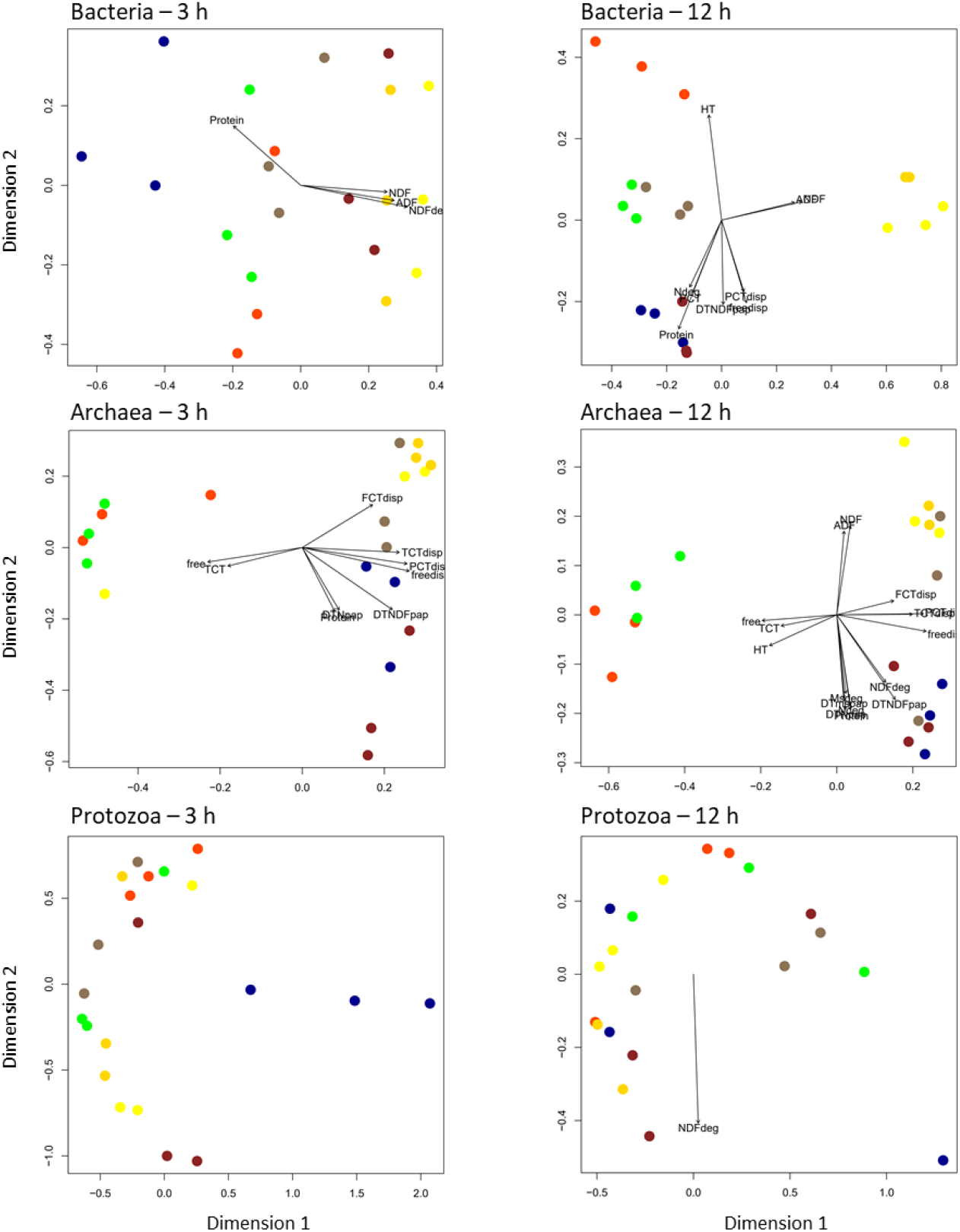
Relationship between chemical composition and degradability in the rumen of tropical tannin-rich plants and their associated microbial communities after 3 and 12 h of incubation. 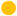= control; 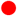 = *Acacia nilotica*; 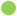 = *Calliandra calothyrsus*; 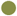 = *Gliricidia sepium*; 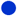 = *Leucaena leucocephala*; 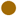 = *Manihot esculenta*; 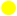 = *Musa spp*. Variables included in the analysis were NDF, ADF and N content; DM, NDF and N degradability (Deg) after 3 or 12 h of incubation; DM, N and NDF theoretical degradability (DT); contents in hydrolysable (HT) and total condensed tannins (TCT) and their fractions (free for free CT, FCT for fibre-bound CT, PCT for protein-bound CT); disappearance (dis) of total condensed tannins and their fractions. Variables significantly correlated are depicted in the graphs.

**Supplementary Figure 3.**
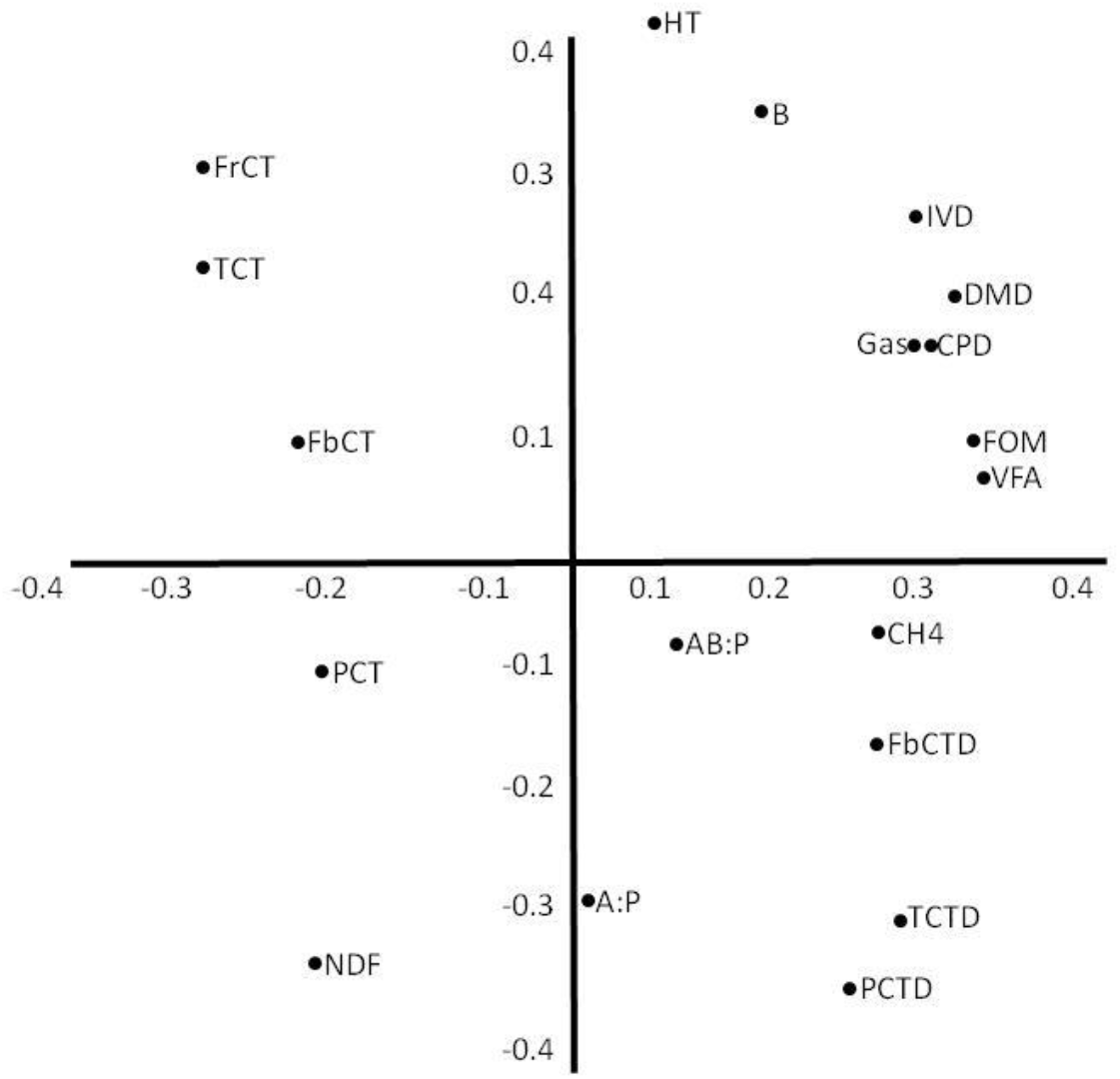
Contribution of variables of tropical tannin-rich plants and variables obtained in vitro and in situ. Abscissa and intercept represent the two main components accounting for 49 and 21% of total variability, respectively. Values in the axes are eigenvalues of the correlation matrix. Observations are average values for each tannin-rich plant (n = 6). Variables of feed characterisation: NDF: neutral detergent fibre; FbCT: fibre-bound condensed tannins; FrCT: free condensed tannins; PCT: protein-bound condensed tannins; TCT: total condensed tannins; HT: hydrolysable tannins; IVD: *in vitro* digestibility. Variables from Exp. 1: DMD: dry matter degradability; CPD: crude protein degradability; FbCTD: fibre-bound condensed tannins disappearance; PCTD: protein-bound condensed tannins disappearance; TCTD: total condensed tannins disappearance. Variables from Exp. 2: VFA: total volatile fatty acid production; A:P: acetate:propionate ratio; AB:P: (acetate + butyrate):propionate ratio; B: butyrate, % total volatile fatty acids; CH4: methane production; Gas: total gas production; FOM: fermented organic matter.

